# Integrated spatial genomics in tissues reveals invariant and cell type dependent nuclear architecture

**DOI:** 10.1101/2021.04.26.441547

**Authors:** Yodai Takei, Shiwei Zheng, Jina Yun, Sheel Shah, Nico Pierson, Jonathan White, Simone Schindler, Carsten Tischbirek, Guo-Cheng Yuan, Long Cai

## Abstract

Nuclear architecture in tissues can arise from cell-type specific organization of nuclear bodies, chromatin states and chromosome structures. However, the lack of genome-wide measurements to interrelate such modalities within single cells limits our overall understanding of nuclear architecture. Here, we demonstrate integrated spatial genomics in the mouse brain cortex, imaging thousands of genomic loci along with RNAs and subnuclear markers simultaneously in individual cells. We revealed chromatin fixed points, combined with cell-type specific organization of nuclear bodies, arrange the interchromosomal organization and radial positioning of chromosomes in diverse cell types. At the sub-megabase level, we uncovered a collection of single-cell chromosome domain structures, including those for the active and inactive X chromosomes. These results advance our understanding of single-cell nuclear architecture in complex tissues.

## Main Text

The three-dimensional (3D) organization of the genome is critical for many cellular processes, from regulating gene expression to establishing cellular identity (*1–3*). Genome organization has been extensively examined using sequencing-based genomics and microscopy approaches (*4, 5*). In particular, chromosome architectures, such as topologically associating domains (TADs) (*6–8*) and high-order chromosomal interactions (*9, 10*), have been revealed by high-throughput genomics approaches such as Hi-C (*11*), genome architecture mapping (GAM) (*9*), and split-pool recognition of interactions by tag extension (SPRITE) (*10*). Moreover, recent progress in chromosome capture methods has enabled the exploration of chromosome structures at the single-cell level (*12–19*). These studies have characterized the variabilities of chromosome structures in single cells derived from various biological samples. Complementary to the genomics approaches, imaging-based approaches such as DNA fluorescence in situ hybridization (FISH) (*20*) can directly obtain 3D chromosome structures from measured loci in single cells without computational reconstructions. Recent multiplexed imaging-based methods (*21–32*), such as sequential DNA FISH (*22*) and in situ genome sequencing (IGS) (*31*), have directly characterized the variabilities of chromosome structures in 3D, even between homologous chromosomes in a cell, and reported TAD-like domain structures in single cells (*23*), which when averaged over populations of cells are consistent with sequencing-based bulk measurements. Furthermore, super-resolution imaging studies (*23, 33, 34*) have shown that architectural proteins such as CTCF and cohesin can play important roles in the single-cell domain structures.

To better understand the principles underlying chromosome organization, it is crucial to integrate chromosome structures with other measurements that capture transcriptional states (*35*), chromatin states (*36*), nuclear bodies (*37, 38*) and radial organization of the nucleus (*39*) in single cells. Single-cell multimodal genomics technologies (*40*) can evaluate chromosome structures together with, for instance, DNA methylome profiling (*41, 42*). However, sequencing-based single-cell multimodal measurements of chromosome structure and the transcriptome in the same cell remain challenging. On the other hand, imaging-based approaches allow direct integration of multimodal measurements including chromosome structures (*22, 23, 25, 27–32*).

We recently established an imaging-based integrated spatial genomics approach that enables the analysis of nuclear organization beyond chromosome structures (*32*). Briefly, we imaged thousands of genomic loci by DNA seqFISH+ along with transcriptional states by RNA seqFISH and subnuclear localization of histone modifications and nuclear bodies by sequential immunofluorescence (IF) in single mouse embryonic stem (ES) cells. We discovered that chromosomes consistently associate with specific nuclear bodies, such as nuclear speckles (*43*) and nucleolus (*44*), across many single cells. We found that individual chromosomes contain unique combinations of fixed loci that are consistently associated with different nuclear bodies and chromatin modifications, suggesting a scaffolding of chromosomes across multiple nuclear bodies and protein globules. Nevertheless, to what extent these principles for nuclear organization extend to diverse cell types, and in a complex and physiologically relevant context such as mammalian tissues, is largely unknown. Imaging-based multimodal measurements of multiple cell types in tissues will give us a great opportunity to dissect cell-type specific features and invariant principles in nuclear organization in the native context.

### Integrated spatial genomics in the brain

To comprehensively investigate the principles of nuclear organization among single cells in a tissue, we analyzed sections of the adult mouse cerebral cortex. Specifically, we applied our integrated spatial genomics approach (*32*) to evaluate 3,660 DNA loci, 76 cellular RNAs, and 8 chromatin marks and nuclear bodies (*45*) (Fig. 1 and fig. S1 and table S1 to S4). This technology provides a powerful multimodal tool to integrate the transcriptional states, chromosome and subnuclear structures, radial nuclear organization, and chromatin and morphological features simultaneously in the same cells in tissues. We chose the mouse brain as a model as it comprises many distinct cell types and has been extensively studied by single cell RNA sequencing (*46–48*) as well as by spatial transcriptomics methods (*49–54*).

**Fig. 1.**
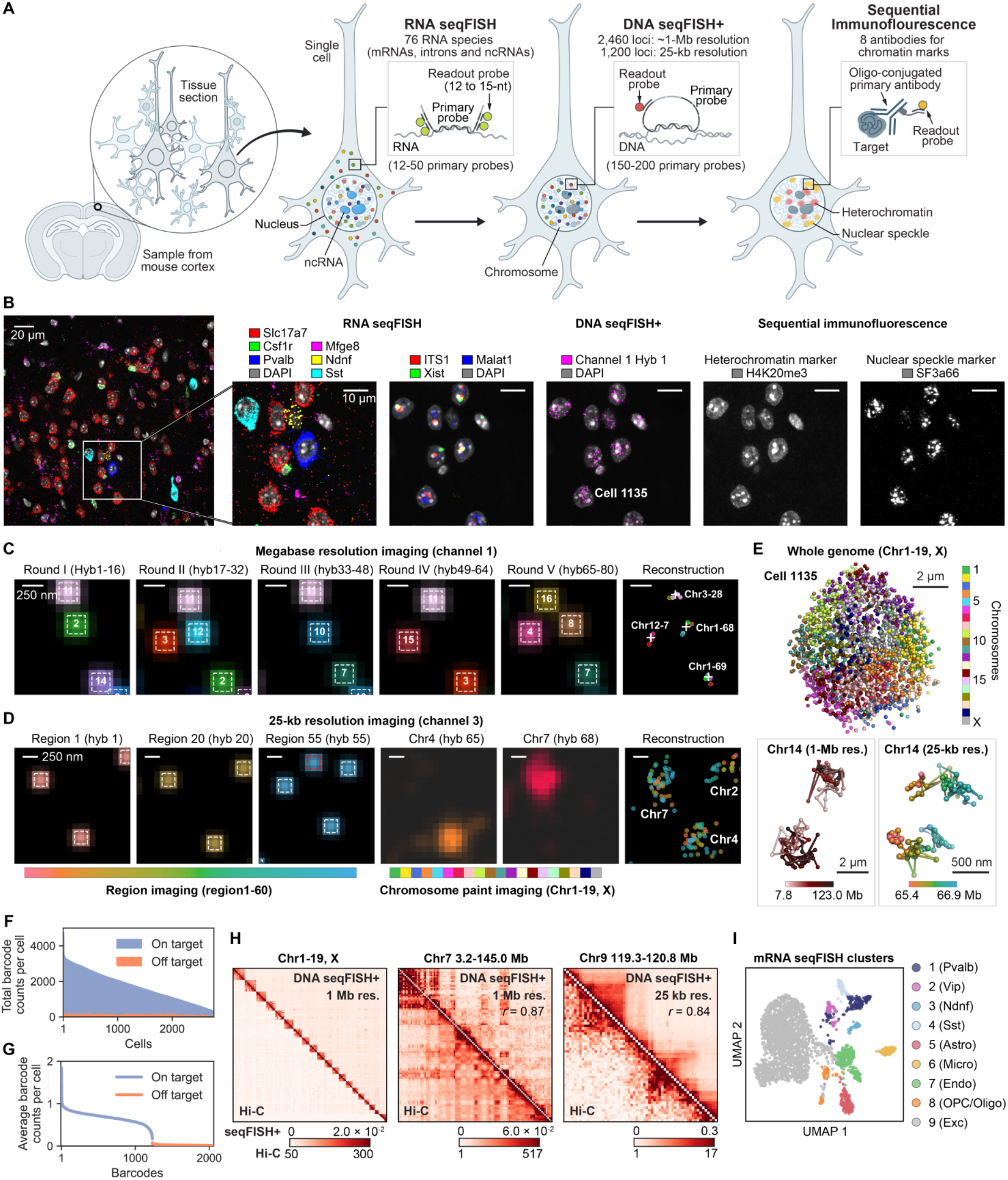
Integrated spatial genomics in the mouse brain. (**A**) Schematic of integrated RNA seqFISH, DNA seqFISH+ and sequential immunofluorescence (IF) measurements in the mouse brain cortex. 76 RNA species targeting introns, mRNAs and noncoding RNAs were imaged first by RNA seqFISH, followed by 2,460 genomic loci imaged at 1-Mb resolution and 1,200 genomics loci at 25-kb resolution by DNA seqFISH+. Finally, 8 antibodies targeting chromatin marks and nuclear bodies were imaged by sequential IF, as well as 5 repetitive elements in the genome by DNA FISH. (**B**) Example RNA seqFISH, DNA seqFISH+ and sequential IF images in a brain slice. Cell type specific RNA expression labels distinct neuronal and glial cell types (e.g. Slc17a7 in excitatory neurons, Mfge8 for astrocytes, Pvalb for inhibitory neurons), along with nuclear body markers, DNA seqFISH+ from 1 round of hybridization, and immunofluorescence signals in the same cells. The images are the maximum intensity z-projected from 1.5 μm z-slices. Scale bars represent 20 μm (left) and 10 μm (right panels). (**C**) Example DNA seqFISH+ images for the 1 zoomed-in view of Cell 1135 through five rounds of barcoding in fluorescent channel 1 decoded with megabase resolution from a single z-section. Images from 16 serial hybridizations are collapsed into a single composite 16-pseudocolor image, which corresponds to one barcoding round. (**D**) The zoomed-in view of Cell 1135 through 60 hybridization rounds targeting adjacent regions at 25-kb resolution followed by 20 hybridization rounds of chromosome painting in fluorescent channel 3. The images are shown with pseudocolors with spots from all z-slices from the nucleus. White boxes on pseudocolor spots indicate identified barcodes in (C) and (D), and the red box indicates a rejected non-specific binding spot in (D). Scale bars represent 250 nm in top and bottom zoomed-in images in (C) and (D). (**E**) 3D reconstruction of a single nucleus. Top, individual chromosomes labeled in different colors. Lower left, reconstruction of chromosome 14 colored based on chromosome coordinates. Lower right, reconstruction of chromosome 14 at from 65.4-66.9 Mb with 25-kb resolution. (**F**) Frequencies of on-and off-target barcodes in fluorescent channel 1 and 2 per cell. On average, 1,617.5 ± 817.1 (median ± standard deviation) on-target barcodes and 32.0 ± 27.8 off-target barcodes are detected per cell (n = 2,762 cells from three biological replicates). (**G**) Average frequencies of individual on-target and off-target barcodes (n = 2,460 barcodes in fluorescent channel 1 and 2) calculated from n = 2,762 cells in (F), demonstrating the accuracy of the DNA seqFISH+. (**H**) Agreement between the normalized spatial proximity maps (probability of pairs of loci within 500 nm in cells for 1-Mb resolution data and within 150 nm in alleles for 25-kb resolution data) from DNA seqFISH+ (upper right) and Hi-C (lower left) (*6, 56*) at different genomic scales with whole chromosomes (left), whole chromosome 7 at 1-Mb resolution (middle), and chromosome 9 region at 25-kb resolution (right) (n = 2,762 cells from three biological replicates). Hi-C data is displayed with 1-Mb and 25-kb bin sizes to compare with 1-Mb and 25-kb resolution DNA seqFISH+ data, respectively. Similar results were obtained for other chromosomes and regions (fig. S2). (**I**) UMAP representation of the cell type clusters determined based on mRNA seqFISH profiles with 2,762 cells from three biological replicates. n = 155, 58, 41, 53, 152, 90, 240, 78, 1,895 cells in each cluster. The transcriptionally defined cell types match with scRNA-seq data (*47*) in fig. S1E.

We examined chromosome structures of the entire genome by using DNA seqFISH+ to image 2,460 loci, at approximately 1-Mb resolution, in 20 chromosomes. These data were collected using a 16 pseudocolor seqFISH+ coding scheme (*32, 53*) in two independent fluorescent channels (Fig. 1C and table S1). In addition, for each of the 20 chromosomes, we examined a local region of at least 1.5 Mb by imaging an additional 1,200 loci at 25-kb resolution (*32*). We collected these data in one fluorescent channel by imaging 60 consecutive loci on each of the 20 chromosomes (Fig. 1D and table S1). In single cells, individual chromosomes formed distinct chromosome territories that have highly variable structures (Fig. 1E). We detected 2,813.0 ± 1,334.0 (median ± standard deviation) spots per cell in total across three fluorescent channels in 2,762 cells from three biological replicates (Fig. 1F, 1G and fig. S1A, S1G and S1H (*55*)). This corresponds to an estimated detection efficiency of at least 38.4 ± 18.2% (median ± standard deviation) in post-mitotic cells with the diploid genome. On the other hand, the false positive spots, as determined by the barcodes unused in the codebook (table S1), were detected at 32.0 ± 27.8 (median ± standard deviation) per cell (Fig. 1F and 1G).

We validated our DNA seqFISH+ data by comparing it with bulk Hi-C data from mouse cortex (*6, 56*) (Fig. 1H and fig. S2A to D). The Hi-C normalized read counts correlated with the mean spatial proximity probability, with a Pearson correlation coefficient of 0.89 and 0.76 at the 1-Mb and 25-kb resolution, respectively (fig. S2A to D). Similarly, Hi-C normalized read counts showed high concordance with DNA seqFISH+ spatial distance at 1-Mb and 25-kb resolution (fig. S2A to D). Furthermore, the DNA seqFISH+ data from the three biological replicates were highly reproducible, with a Pearson correlation coefficient of 0.95-0.97 for 1-Mb and 25-kb resolution data (fig. S2E and S2F). These results demonstrate the robustness of DNA seqFISH+ to map 3D chromosome structures in tissue samples.

We clustered the RNA seqFISH data and obtained 9 major cell type clusters within the cerebral cortex, based on the gene expression of known markers (fig. S1C and S1D), shown in the UMAP representation (Fig. 1I). These 9 clusters matched well with cell types identified from single-cell RNA sequencing (*47*) (fig. S1E). The majority of the cells were excitatory neurons in cluster 9 expressing Slc17a7 (Fig. 1I and fig. S1D). We also observed four subclasses of inhibitory neurons expressing Pvalb, Vip, Ndnf, or Sst, three types of glial cells (astrocytes expressing Mfge8, microglia expressing Csf1r and oligodendrocyte precursor cells and oligodendrocytes expressing Olig1), and endothelial cells expressing Cldn5 (fig. S1D). We observed a similar localization accuracy of the FISH spots (fig. S1F) and number of decoded DNA loci (fig. S1H) across different cell type clusters.

In addition to the genome and RNA imaging, we used sequential IF to detect 6 histone modifications or variants (H3K4me2, H3K27me2, H3K27me3, H3K9me3, H4K20me3, mH2A1), nuclear speckles (SF3a66) and the methyl CpG binding protein MeCP2 (Fig. 2A and fig. S1K, S1L and S3A and table S3). We incubated tissue sections with oligonucleotide-conjugated primary antibodies (*57, 58*) prior to the RNA seqFISH steps and sequentially read out the antibody signals with fluorescently labeled probes (*32*), allowing the multiplexed detection of primary antibodies from the same single cells in tissues. The protocol previously used for cell culture experiments (*32*) was optimized for tissue sections to preserve the nuclear structure and accurately align the IF, RNA seqFISH and DNA seqFISH+ data over a total of 125 rounds of hybridizations and imaging on an automated confocal microscope (*45*) (fig. S1I and S1J). Additionally, we performed sequential RNA and DNA FISH to detect 3 non-coding RNA that mark the inactive X chromosome (Xi, Xist), the nucleolus (ITS1), nuclear speckles (Malat1) (*43, 44, 59*) as well as 5 repetitive regions (LINE1, SINEB1, MajSat, MinSat, Telomere) that relate to nuclear organization (*60–62*) (fig. S1K, S1L and S3A).

**Fig. 2.**
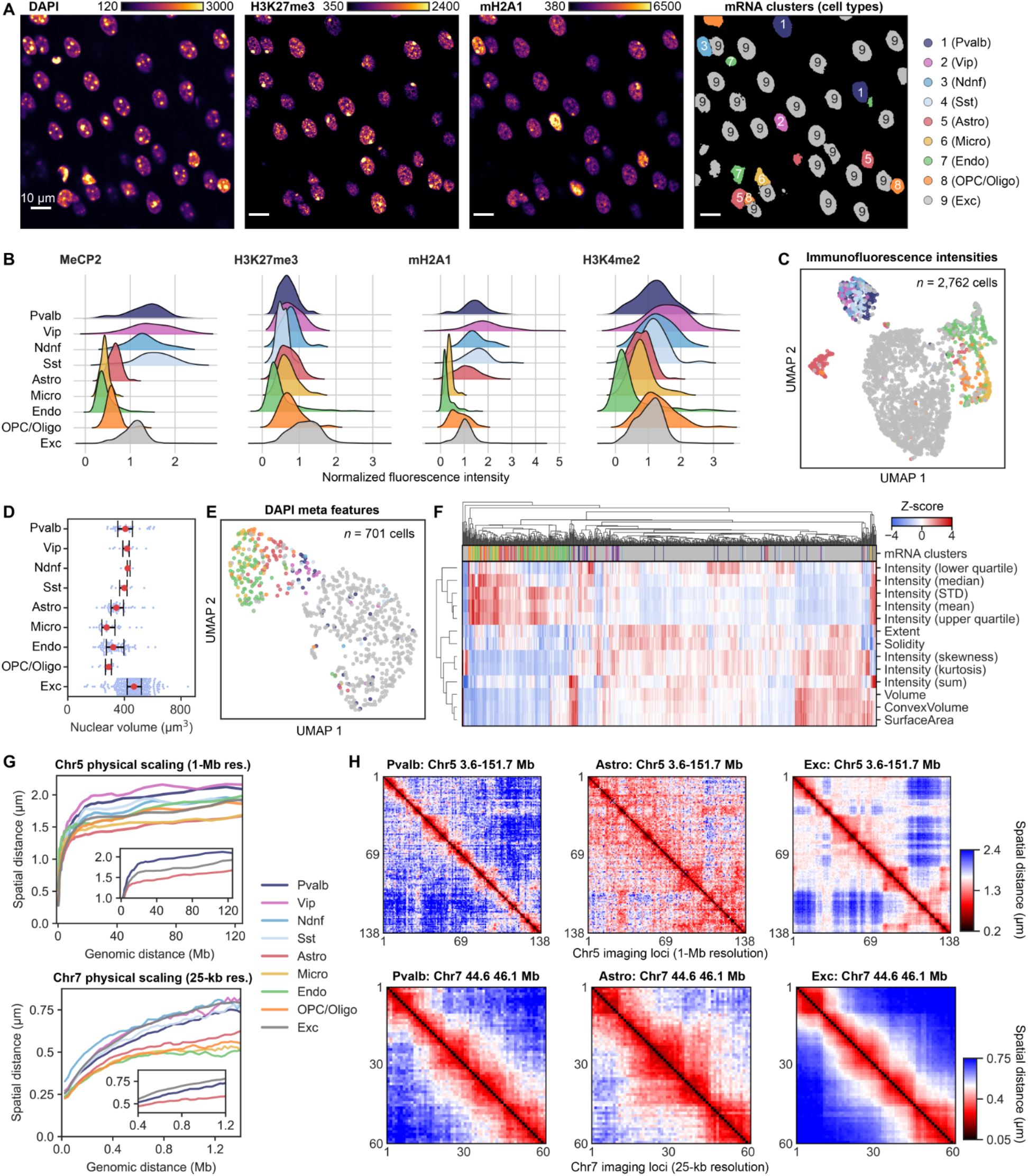
Nuclear morphology, global chromatin states and chromosome scaling are cell type dependent. (**A**) DAPI, H3K27me3, and mH2A1 staining from a single z-section super-imposed on the transcriptionally defined cell clusters. Color bars represent fluorescence intensity (a.u.). Scale bars, 10 μm. (**B**) Kernel density estimations of normalized IF intensities show distinct cell type dependent distributions of histone modification and chromatin marker levels. Additional antibody intensities are shown in fig. S3B. (**C**) UMAP representation shows separation of excitatory, inhibitory and glial cells based on the overall intensities for 8 IF markers in single cells. Cells are colored by their transcriptional cell types in (A). 2,762 cells from three biological replicates were used in (B) and (C). (**D**) Characterization of the nuclear volume across the 9 major cell types. The red dots represent median, and whiskers represent the interquartile range. (**E**) UMAP representation of cells based on DAPI meta features used in (F). Cells are colored by their transcriptional cell types in (A). (**F**) Hierarchical clustering of the DAPI meta features compared to the RNA seqFISH clusters shown on top. 701 cells in the center z-sections from three biological replicates were used in (D)-(F). (**G**) Physical distance as a function of genomic distance across transcriptional defined cell types at the 1-Mb resolution for Chr5 (top) and at 25-kb resolution for Chr7 (bottom). Inset shows Pvalb inhibitory neurons, astrocytes and excitatory neurons. Additional scaling relationships are shown in fig. S3C to E. (**H**) Spatial distance between pairs of intra-chromosomal loci within cells by DNA seqFISH+ in different cell types at 1-Mb resolution (top row, quartile spatial distance within cells) and at 25-kb resolution (bottom row, median spatial distance within alleles). 2,762 cells were used for (G) and 155 cells for Pvalb, 152 cells for astrocytes, and 1,895 cells for excitatory neurons were used in (H) from three biological replicates.

The spatial overlap between individual antibody, RNA and DNA markers of nuclear bodies or subnuclear compartments was consistent with previously observed subnuclear localization patterns (*37, 43, 44, 59–64*). For example, individual cells displayed colocalization of nuclear speckle components (Malat1 and SF3a66), inactive X chromosome (Xi) components (Xist, mH2A1, H3K27me3, LINE1) and heterochromatin components (DAPI, MajSat, H3K9me3, H4K20me3, MeCP2) (fig. S1K and S1L). We also observed minimum spatial overlap between different nuclear bodies such as nuclear speckles (Malat1 and SF3a66) and the nucleolus (ITS1), as well as between euchromatin- (SINEB1) and heterochromatin- (LINE1) enriched chromosomal regions, as expected (*61*) (fig. S1K and S1L). Taken together, these results demonstrate that our integrated spatial genomics approach allows us to explore nuclear organization with unprecedented detail at the RNA, DNA and protein level across diverse cell types in intact tissues.

### Distinct nuclear features across cell types in the brain

We first examined the differences in global histone modifications and variants across cell types in the mouse cortex to understand cell-type specific global chromatin states. The overall intensities of IF markers were highly variable across single cells (Fig. 2A and fig. S3A). The 9 major cell types displayed clear differences in the global levels of both repressive marks (MeCP2, H3K27me3 and mH2A1) and an active mark (H3K4me2) (Fig. 2B). Clustering of single cells using the multiplexed IF data was able to distinguish the same 9 cell types as identified by RNA seqFISH (Fig. 2C), supporting a strong correlation between global chromatin states and transcriptional states. In particular, we observed a relative enrichment of mH2A1 in inhibitory neurons and astrocytes, H3K4me2 in inhibitory neurons, oligodendrocyte precursor cells and oligodendrocytes, and H3K27me3 in excitatory neurons (Fig. 2A and 2B and fig. S3A and S3B). In addition, MeCP2 was enriched in inhibitory and excitatory neurons while lower and almost undetectable levels of intensities were observed in astrocytes and microglia, respectively (Fig. 2A and 2B and fig. S3A). This observation agrees with the previously reported MeCP2 immunostaining intensity in neurons, astrocytes and microglia in the mouse cortex (*65*).

Interestingly, even the DAPI features alone were sufficient to separate the major cell types in the cortical areas of the mouse brain using both UMAP and hierarchical clustering (Fig. 2A and 2D to F). We used a subset of 701 cells for nuclei that were fully covered in the brain section. Compared to glial cells, neurons typically had larger nuclear volumes (Fig. 2D and 2F) and lower DAPI intensities per voxel (e.g. mean and median) (Fig. 2F), consistent with the same DNA content in both cell types. In addition, among neuronal cell types, we also observed a larger nuclear volume in excitatory neurons compared to those in inhibitory neuron subtypes (Fig. 2D and 2F). These results are consistent with the observation that nuclear morphological features are often sufficiently distinct in mammalian tissues (*66*) to determine major cell types by visual inspection of the images.

Lastly, we examined the spatial distance between pairs of intra-chromosomal loci to understand cell-type specific chromosomal scaling in the nucleus. Although previous imaging studies in cell culture and embryos showed differences in the chromosomal scaling of spatial distance as a function of genomic distance (*22, 31, 32*), it remains unclear how the scaling principles operate across different cell types within the same tissues. We found that the relationship between spatial versus genomic distance scaling was distinct in different cell types, in both 1-Mb and 25-kb resolution data (Fig. 2G and 2H and fig. S3C to G). For example, loci tens of megabases apart typically displayed a larger spatial separation in inhibitory neurons compared to excitatory neurons and glial cells (Fig. 2G and fig. S3C), which cannot be simply explained by nuclear size differences as inhibitory neurons typically had smaller nuclear sizes than excitatory neurons (Fig. 2D). In contrast, in the 25-kb resolution data at the targeted Mb regions, the scaling relationship differed depending on the chromosomal regions and cell types (Fig. 2G and fig. S3D). For example, the targeted regions in chromosome 7 and chromosome 17 are more dispersed in neurons compared to glial cells, whereas other targeted regions in chromosome 5 and chromosome 18 have almost identical scaling relationships among neurons and glial cells. Overall, regions with higher gene density tend to have less compact spatial organization (fig. S3E), possibly owing to different underlying epigenetic states (*32, 67*). To gain a more integrative picture of nuclear organization, we need to examine the interactions between DNA and nuclear bodies beyond characterizing individual components.

### DNA loci display unique and shared chromatin profiles across different cell types

To characterize the spatial association between DNA loci, chromatin marks and nuclear bodies, we calculated the fraction of time that each DNA locus is associated with each chromatin mark. Because many chromatin marks form discrete regions within the nucleus and IF images are diffraction limited, we determine the fraction of time each DNA locus is within 300 nm from the exterior of each mark (*32, 55*) (Fig. 3A and 3B and fig. S4A). This imaging-based approach of computing “chromatin profiles” demonstrated a high correlation with sequencing-based bulk measurements such as ChIP-seq and SPRITE at 1-Mb resolution in mouse ES cells (*32*).

**Fig. 3.**
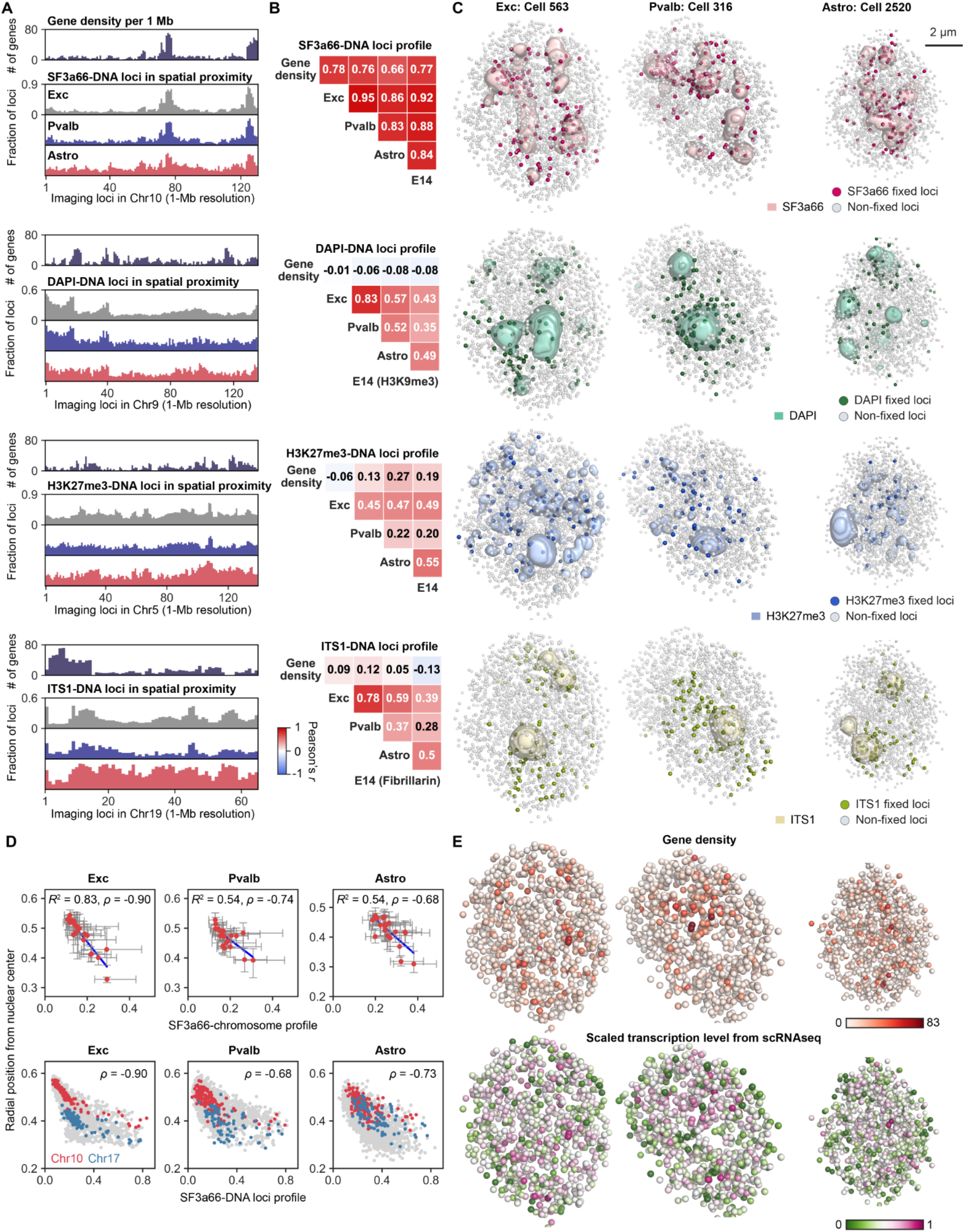
Chromatin profiles and fixed points across cell types. (**A**) Cell type specific chromatin profiles, the fraction of loci found within 300 nm of SF3a66, DAPI, H3K27me3 and ITS1 exteriors. Gene density within 1-Mb bin size is shown for comparison. (**B**) Pearson correlation between chromatin profiles of different cell types and gene density for autosomes (n = 2,340 loci). Correlation with the chromatin profiles from E14 mouse ES cells (*32*) are shown for comparison. (**C**) Representative 3D images for fixed loci and IF markers. For IF marks, pixels with intensity z-score values above 2 for each IF mark were shown. Fixed loci were determined by z-score above 2 from loci in autosomes. (**D**) Nuclear speckle association scores are correlated with radial nuclear positioning for 20 chromosomes (top panels) and 2,460 individual loci (bottom) such that nuclear speckle associated chromosomes or loci position at more nuclear interior. The red dots represent median for individual chromosomes and error bars represent interquartile ranges (top). The coefficient of determination for linear regression (top) and Spearman correlation coefficients (top and bottom) are shown. (**E**) In single cells, loci with higher gene density (top) and higher 1-Mb resolution transcription levels obtained from cell-type-specific expression profiles by scRNA-seq (*47*) (bottom) appear around nuclear speckles, and are not necessarily radially distributed from nuclear interior to exterior. The same cells are shown as in (C) with 2 μm cross sections. n = 1,895, 155, 152 cells for excitatory neurons, Pvalb inhibitory neurons, and astrocytes from three biological replicates in (A), (B) and (D).

Whereas some chromatin profiles were highly concordant across cell types, others showed specific patterns that varied between cell types (Fig. 3A and 3B and fig. S4A). The DNA loci associated with nuclear speckle markers (SF3a66 and Malat1) were highly correlated among different cell types at 1-Mb resolution, with a Pearson correlation coefficient of >=0.83 even including mouse ES cells (Fig. 3B). This highly conserved DNA-nuclear speckle spatial association has been reported recently with various cell lines using TSA-seq (*68*). Interestingly, the chromatin profiles for nuclear speckles were highly correlated with gene densities at 1-Mb resolution (Fig. 3A and 3B and fig. S4A), suggesting a robust relationship between spatial genome organization around nuclear speckles and underlying genomic sequences.

On the other hand, the associations between DNA loci and DAPI-rich constitutive heterochromatin showed lower correlation between neurons and astrocytes (Fig. 3A and 3B), even though the associations were typically enriched in the centromere proximal genomic loci in all cell types (Fig. 3A and fig. S4A and S4B). Furthermore, the relationships between DNA loci and H3K27me3 were more distinct across cell types (Fig. 3A) and showed lower correlation across cell types in the brain as well as with mouse ES cells (Fig. 3B), reflecting the underlying differences in DNA loci-histone modification globule associations across cell types.

We observed similar cell-type-dependent association of chromosomes with the nucleolus in the mouse brain cortex (Fig. 3A). The spatial proximities between ITS1 non-coding RNA and DNA loci in Pvalb inhibitory neurons and astrocytes displayed a relatively low correlation, with a Pearson correlation coefficient of 0.37 (Fig. 3B). We observed some cells in which the 45S ribosomal DNA (rDNA)-containing chromosomes 15 and 19 (*69*) were not in physical proximity to the nucleoli but were close to DAPI-rich heterochromatin regions (fig. S4C), possibly due to rDNA silencing (*70*). This rDNA silencing could lead to the cell-type specific nucleolar association of genomic loci (Fig. 3A and 3B and fig. S4A). To confirm this, we performed imaging of rDNA loci by DNA FISH and found that different fractions of rDNA loci were associated with the nucleolus and DAPI-rich heterochromatin regions among neurons, astrocytes, mouse ES cells and cultured fibroblasts (fig. S4D to F), lending support to the notion that nucleolar organizer regions can be stably silenced (*70*) in a tissue-specific fashion.

### Cell-type specific fixed loci anchor chromosomes to nuclear bodies in single cells

To further gain insight of single-cell nuclear architecture across cell types, we defined DNA loci that were consistently associated with a particular chromatin mark or nuclear body in each cell type to be “fixed loci” (*45*) (Fig. 3C). We previously observed that these fixed loci for each IF marker consistently appear on the exterior of the respective marker in single mouse ES cells (*32*). In the mouse brain, we observed similar associations of the fixed loci with the exterior of nuclear bodies and chromatin marks in single cells (Fig. 3C), despite the differences in the morphological features and arrangement of nuclear bodies in individual neuronal and glial cells. As examples, fixed loci for SF3a66, DAPI, and H3K27me3 are consistently observed on the exterior of nuclear speckles, heterochromatin bodies and H3K27me3 globules in single neurons and glial cells (Fig. 3C).

Fixed loci for different chromatin marks are distributed across the genome such that each of the chromosomes has distinct patterns of fixed loci in each cell type (Fig. 4A to D and fig. S5A). These fixed loci can constrain the nuclear organization of chromosomes by their association to the nuclear bodies or chromatin marks in individual nuclei (Fig. 4C and 4D and fig. S5A). For example, chromosomes 7 and 17 have fixed loci for nuclear speckles (SF3a66) and heterochromatic bodies (DAPI) in excitatory neurons, inhibitory neurons and astrocytes (Fig. 4D), and both chromosomes straddled these nuclear bodies in single cells of all three cell types. Similarly, chromosome 8 has nuclear speckles and H3K27me3 fixed points in all three cell types, and spanned those nuclear globules in single cells. A small number of loci were associated with two nuclear bodies (orange dots) and were observed near both nuclear bodies in single cells (Fig. 4C and 4D). These features were observed across different cell types for other chromosomes (fig. S5A). Thus, despite the differences in the arrangement of nuclear bodies in individual cells and the different fixed point patterns on the chromosomes in each cell type (Fig. 4D and fig. S5A), the association between DNA loci and nuclear bodies are consistent across single cells of each cell type. These findings in the mouse brain extend our previous work in mouse ES cells (*32*) to show that chromatin fixed loci serve as organizational invariants in the nuclei of single cells across cell types, despite the highly variable appearance of individual chromosomes and nuclear bodies in individual cells.

**Fig. 4.**
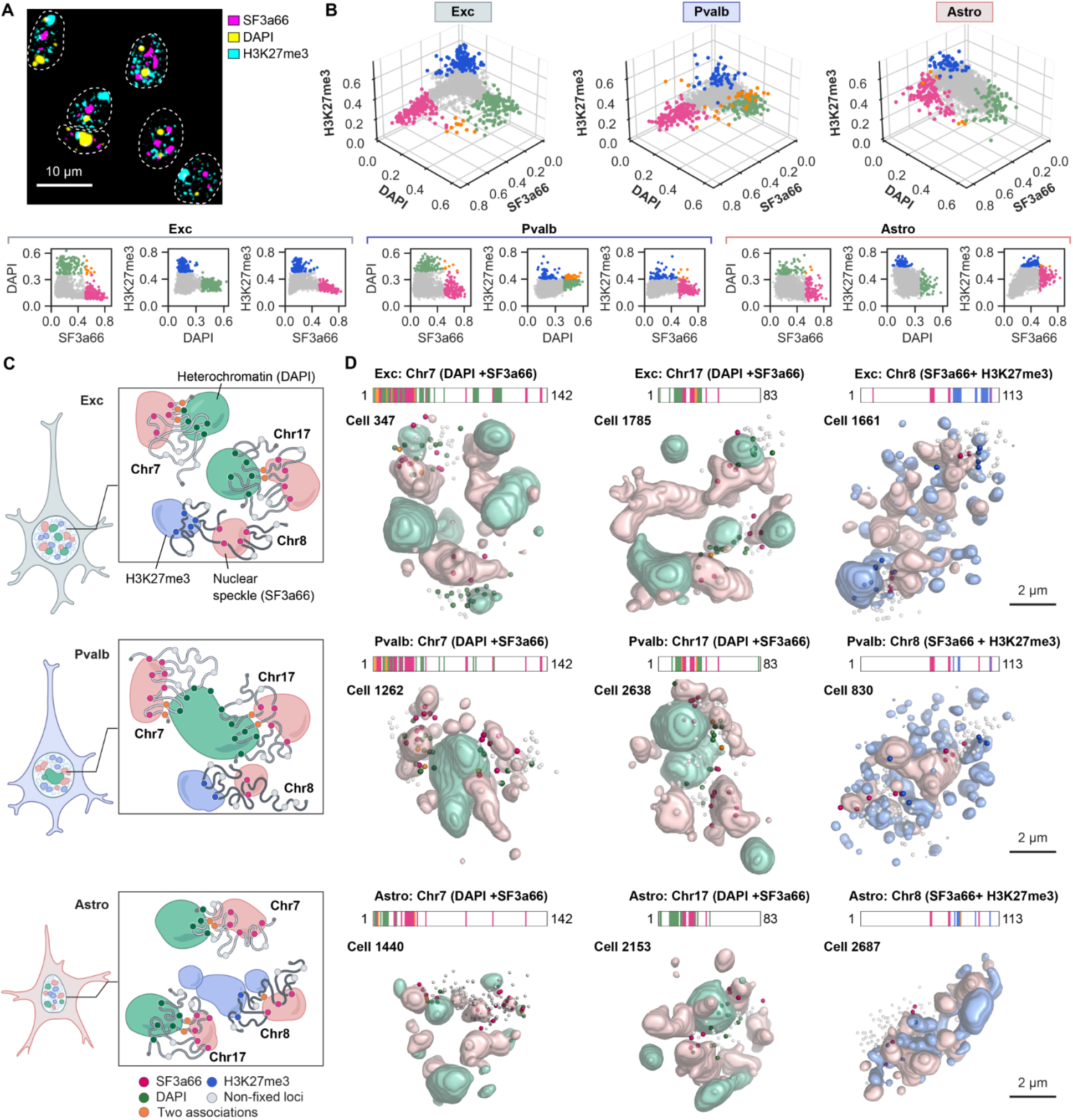
Fixed points straddle across nuclear bodies. (**A**) Heterochromatin stained by DAPI, nuclear speckles marked by SF3a66 and H3K27me3 globules are minimally overlapping in individual nuclei (white dashed lines) in the mouse cortex section. (**B**) Joint profiles for the three markers for each genomic locus in autosomes (n = 2,340 loci) in excitatory neurons (left), Pvalb inhibitory neurons (middle), and astrocytes (right) computed from the population of cells. Loci with high association scores in each of the marks are highlighted in the respective color (green for DAPI, magenta for SF3a66, and blue for H3K27me3). Loci with high association scores for two markers are labeled in orange. Bottom panel shows pairwise association scores for all the loci. n = 1,895, 155, and 152 cells for excitatory neurons, Pvalb inhibitory neurons, and astrocytes from three biological replicates. (**C**) Illustration showing chromosome 7 and 17 with fixed loci for SF3a66 and DAPI and chromosome 8 with fixed loci for SF3a66 and H3K27me3 across cell types (Exc, excitatory neurons; Pvalb, Pvalb inhibitory neurons; Astro, astrocytes). Fixed loci associated with two markers (both SF3a66 and DAPI in Chr7 and Chr17, and both SF3a66 and H3K27me3 in Chr8) are shown with orange color. (**D**) Representative 3D images of individual chromosomes (Chr7, 8, or 17) and markers (SF3a66, H3K9me3 or H3K27me3) showing each chromosome has cell-type specific fixed points and span the corresponding nuclear bodies in single cells. The same coloring as (C). For other chromosomes and cells, see fig. S5A.

### Cell-type specific nuclear bodies determine inter-chromosomal proximity and radial positioning

The cell-type specific inter-chromosomal interaction and radial positioning of chromosomes in the nucleus were previously characterized by chromosome paint (*71–73*) and single-cell chromosome conformation capture (*18, 19*). However, it remains unclear what determines the distinct chromosomal features across cell types. Similarly, although the arrangement of nuclear bodies, such as heterochromatin bodies and nucleoli, has been shown to be cell-type specific as well as dynamic even within a cell type during development (*61, 74*), it remains unclear how those differences in nuclear body arrangements relate to 3D chromosome organization at the single-cell resolution. Thus, we examined our integrated spatial multi-modal datasets to test the hypothesis that nuclear body organization and the fixed point association in different cell types accounts for the cell-type specific inter-chromosomal interaction and radial chromosomal positioning.

We first characterized the nuclear bodies in each cell type (Fig. 5A to C and fig. S5B). Compared to excitatory neurons and astrocytes, Pvalb inhibitory neurons had fewer but larger heterochromatic bodies and nucleoli (Fig. 5B and fig. S5B). Heterochromatic bodies were often localized close to the center of the nucleus in Pvalb inhibitory neurons, but more distributed in other cell types (Fig. 5C). Nuclear speckles appeared to be more dispersed in cells with preferred localization in the nuclear interior (Fig. 5A and 5C). Furthermore, H3K27me3 globules tend to localize more at the nuclear interior in astrocytes as compared to neurons (Fig. 5A and 5C).

**Fig. 5.**
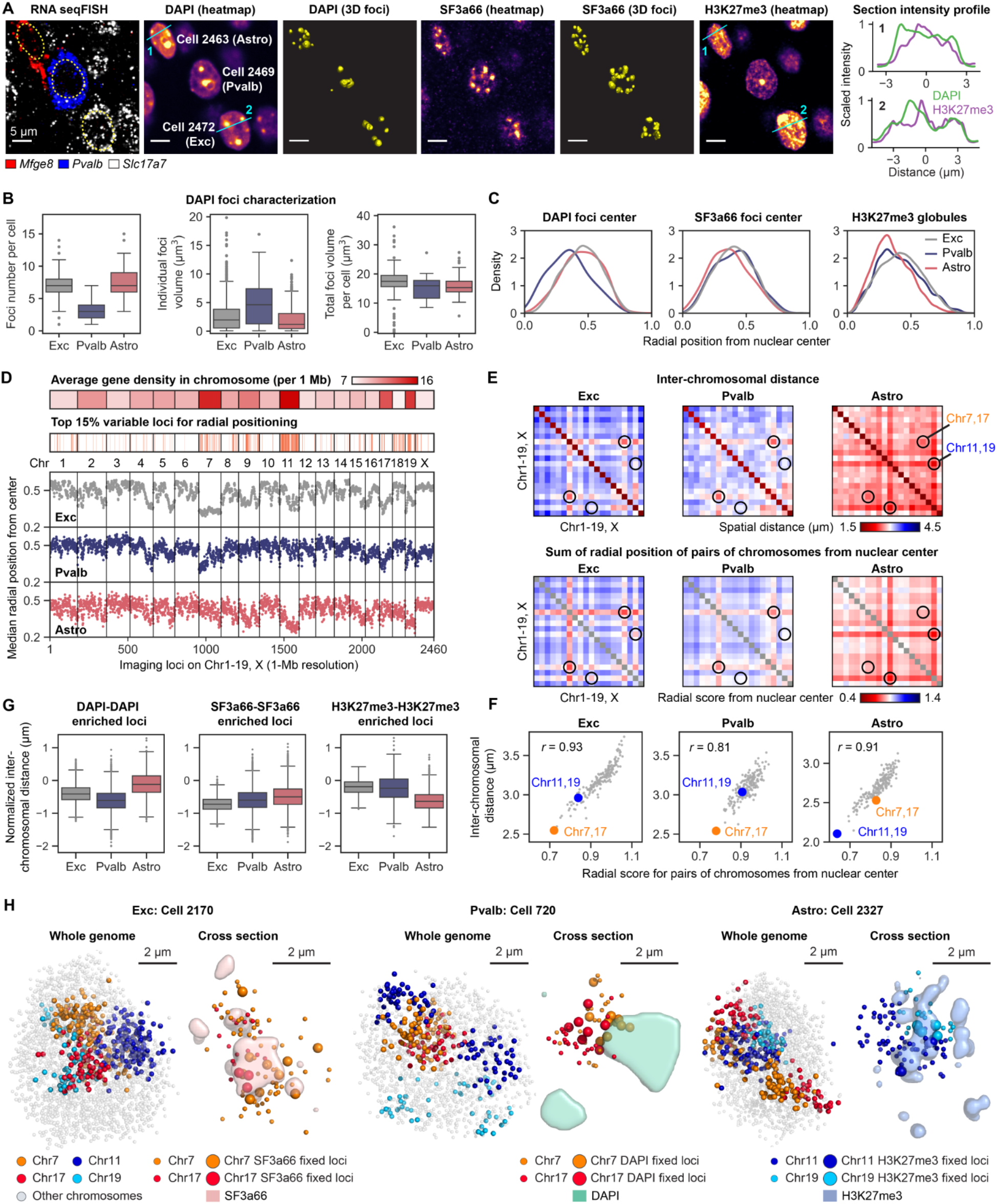
Fixed points and nuclear bodies organize nuclear architecture. (**A**) DAPI, SF3a66 and H3K27me3 staining from a single z-section with transcriptionally defined cell type annotation. Scale bars, 5 μm. Cross-section intensity profiles for DAPI and H3K27me3 for the astrocyte (line 1) and excitatory neuron (line 2) are shown (right), representing the depletion of H3K27me3 signals from the nuclear periphery in astrocytes. (**B**) Characterization of DAPI foci features across 3 major cell types. The boxplots represent the median, interquartile ranges, whiskers within 1.5 times the interquartile range, and outliers. (**C**) Kernel density estimation for the radial positioning of identified nuclear foci centers (DAPI and SF3a66) or Xist-independent H3K27me3 globules calculated by intensity z-score above 2 in each cell for different cell types. n = 460, 33, and 48 cells for excitatory neurons, Pvalb inhibitory neurons, and astrocytes in the center fields of view from three biological replicates in (B) and (C). (**D**) Median radial positioning from the nuclear center for all 2,460 loci imaged at 1-Mb resolution for three major cell types. Gene density (top panel) and the top 15% variable loci in terms of radial positioning among the three cell types (middle panel) are shown. (**E**) Comparison of mean pairwise distance between chromosomes (top panels) and the radial position scores for pairs of chromosomes (bottom panels) for the three major cell types. Chromosomes near the nuclear interior also tend to be spatially closer to other chromosomes. (**F**) Scatter plot of mean pairwise distance between chromosomes and the sum of radial position scores for pairs of chromosomes shown in (E). (**G**) Normalized inter-chromosomal distance between pairs of fixed loci enriched in association with specific nuclear bodies and chromatin marks. Inter-chromosomal pairs of fixed loci for the same marker are generally spatially proximal compared to the other inter-chromosome pairs. In particular, DAPI enriched loci are spatially proximal in Pvalb inhibitory neurons while H3K27me3 enriched loci are closer to each other in astrocytes. The distances are normalized by subtracting an averaged inter-chromosomal spatial distance from all inter-chromosomal pairs of loci in each cell type. The boxplots represent the median, interquartile ranges, whiskers within 1.5 times the interquartile range, and outliers. n = 1,895, 155, and 152 cells for excitatory neurons, Pvalb inhibitory neurons, and astrocytes from three biological replicates in (D)-(G). (**H**) Chromosomes 7 and 17 are closer to the nuclear interior in neurons, shown with the fixed points on nuclear speckles (excitatory neuron, left) and heterochromatin (Pvalb inhibitory neuron, middle). Chromosomes 11 and 19 are closer to the nuclear interior in the astrocyte with the fixed points on H3K27me3 globules (right). 2 μm cross sections are shown for the excitatory and Pvalb inhibitory neuron and a 1 μm cross section is shown for the astrocyte for visual clarity.

Next, we characterized radial positioning of individual chromosomes and individual loci from nuclear interior to exterior, and we observed their changes across cell types (Fig. 5D and fig. S5C and S5D). Consistent with previous reports (*71–73*), we observed the correlation between radial positioning and gene density as well as chromosome size across cell types (fig. S5E and S5F). Interestingly, gene-dense chromosomes (e.g. Chr7, 11, 17 and 19) tend to have different radial positions across different cell types (Fig. 5D and fig. S5C and S5F), which was also observed during post-natal brain development (*19*). For example, gene dense chromosome 7 tends to localize close to the nuclear center in excitatory neurons, but not in glial cells (Fig. 5D and fig. S5C). At the same time, we observed a cell-type dependence in the average inter-chromosome spatial distances (Fig. 5E, top). Notably, those pairwise inter-chromosome distance maps agree with averaged radial positioning of pairs of chromosomes, such that chromosomes in the nuclear interior tend to be spatially close to each other (Fig. 5E bottom, and 5F and fig. S5G). These features were observed across cell types, suggesting a common principle in chromosome organization.

We further investigated how the cell-type dependent changes in nuclear bodies, chromosome radial positioning and inter-chromosome arrangement are connected in single cells. We observed that pairs of inter-chromosomal loci assigned as fixed points at nuclear bodies tend to be spatially closer to each other than the other non-fixed point pairs in neurons and astrocytes (Fig. 5G). These fixed points can influence the cell type-specific arrangement of the chromosomes. For example, chromosomes 11 and 19 have many H3K27me3 fixed points in astrocytes (fig. S5A) and H3K27me3 globules tend to be in the interior of the astrocyte nuclei (Fig. 5C). Thus, chromosomes 11 and 19 tend to be observed near the interior and interact with other chromosomes in astrocytes (Fig. 5E and F), but less in neurons. Similarly, chromosome 17 contains many heterochromatin fixed points in neurons (Fig. 4D) and heterochromatic bodies tend to be in the nuclear interior in Pvalb inhibitory neurons (Fig. 5C). Consequently, chromosome 17 tends to be radially positioned near the nuclear interior (Fig. 5D). In addition, because chromosomes 7 and 17 are gene dense and contain many fixed points to nuclear speckles which tends to be positioned in the nuclear interior, both chromosomes are observed in the nuclear interior of neurons in bulk and single cells (Fig. 5D to F and 5H). Thus, the complexity in the global organization of chromosomes in diverse single cells in the brain, such as the cell type-dependence in radial positioning and inter-chromosomal distances, can be dimensionally reduced and captured in the different nuclear body arrangements and fixed point chromatin profiles.

Lastly, we compared the population-averaged and single-cell picture of radial organization of chromosomes and nuclear bodies. At the population-averaged level, the radial positioning of chromosomes and genomic loci to nuclear interior appear to be correlated with nuclear speckle association (Fig. 3D), consistent with other bulk analysis (*75*). However, visualization of the genomic loci as a function of gene density or expression levels in single cells shows that the radial positioning effect is highly variable in each nucleus (Fig. 3E). Thus, while speckle associations for gene dense regions are highly consistent across cells (Fig. 3A and 3B), because the arrangement of nuclear speckles is heterogeneous in single cells with a weak propensity for nuclear center (Fig. 5C), the positioning of the gene-dense speckle associated loci do not show strong radial gradient from nuclear interior to nuclear exterior in single cells, which directly supports the refined model of radial nuclear organization (*75*).

### Domain boundaries are variable in single cells

The high-resolution DNA seqFISH+ data covering the genomic regions at 25-kb resolution enables us to further examine the domain organization of chromosomes in single cells at sub-megabase scales. Analyses based on bulk-averaged chromosome conformation capture data at the sub-megabase resolution suggest that chromosomes are organized into topologically associating domains (TADs) with clear insulation boundaries (*6–8*). The TADs appear largely unchanged across species (*6, 76*) and their boundaries preferentially reside at CCCTC-binding factor (CTCF)- and cohesin-binding sites (*6, 23*). Single-cell chromosome conformation capture measurements confirmed preferential contacts within TADs in single cells (*12*). In addition, imaging experiments confirmed that TAD-like domain structures exist in single cells and depletion of cohesin resulted in shifting boundaries of the examined TADs stochastically across single cells (*23*). However, it is unexplored how single chromosome domain structures are organized across genomic regions with different bulk TAD characteristics.

To systematically investigate single chromosome domain structures, we first determined whether there are subpopulations of chromosomes with distinct configurations that differ from the bulk averaged configurations in excitatory neurons. We clustered the single chromosome pairwise spatial distance data with 25-kb resolution using principal component analysis (PCA) and visualized by UMAP (Fig. 6A to C). The chromosome 3 region (7.7-9.3 Mb) displayed three domains in the bulk data (Fig. 6D, top). However, analysis of subpopulation of chromosomes that were clustered together (Fig. 6C) showed multiple configurations with different pairwise spatial associations and domain boundaries (Fig. 6D, bottom, and 6E, top, and fig. S6A). We further examined single chromosome structures in each structural cluster, and found that instead of all three major domains being present in single cells, in many cells, only a subset of the domains appeared (Fig. 6E, bottom). In the bulk measurements, domain boundaries corresponded to CTCF and cohesin (RAD21, cohesin subunit) binding sites (*77*) (Fig. 6D), but chromosomes in single cells appeared to stochastically form domains from a subset of these sites (Fig. 6E, bottom). These structures can be observed in single chromosome pairwise spatial distance maps of genomic loci as well as direct visualization of the chromosomal regions (Fig. 6E, bottom). In addition, even regions that did not show clear domain structures in the bulk averaged data contained domain-like structures at the single cell level. For example, chromosome 7 (44.6-46.1 Mb) and chromosome 11 (97.4-98.9 Mb) regions contained several domains in single cells even when no clear domains were visible in the bulk level at the examined spatial scale (fig. S6B to E). Furthermore, there were chromosomal regions that exhibited more deterministic boundaries, such as chromosome 6 (49.4-50.9 Mb) region (fig. S7A and S7B). However, single chromosome subpopulations showed heterogeneity in the spatial proximity of inter-domain organization, as seen by the off-diagonal elements (fig. S7B, top and middle) and the 3D reconstruction of the chromosome structures (fig. S7B, bottom). Similarly, the histone gene cluster in chromosome 13 (21.5-23.8 Mb), known to form a higher-order chromosome structure in a population of cells (*10*), showed heterogeneous higher-order chromosome organization in individual cells (fig. S7C and S7D).

**Fig. 6.**
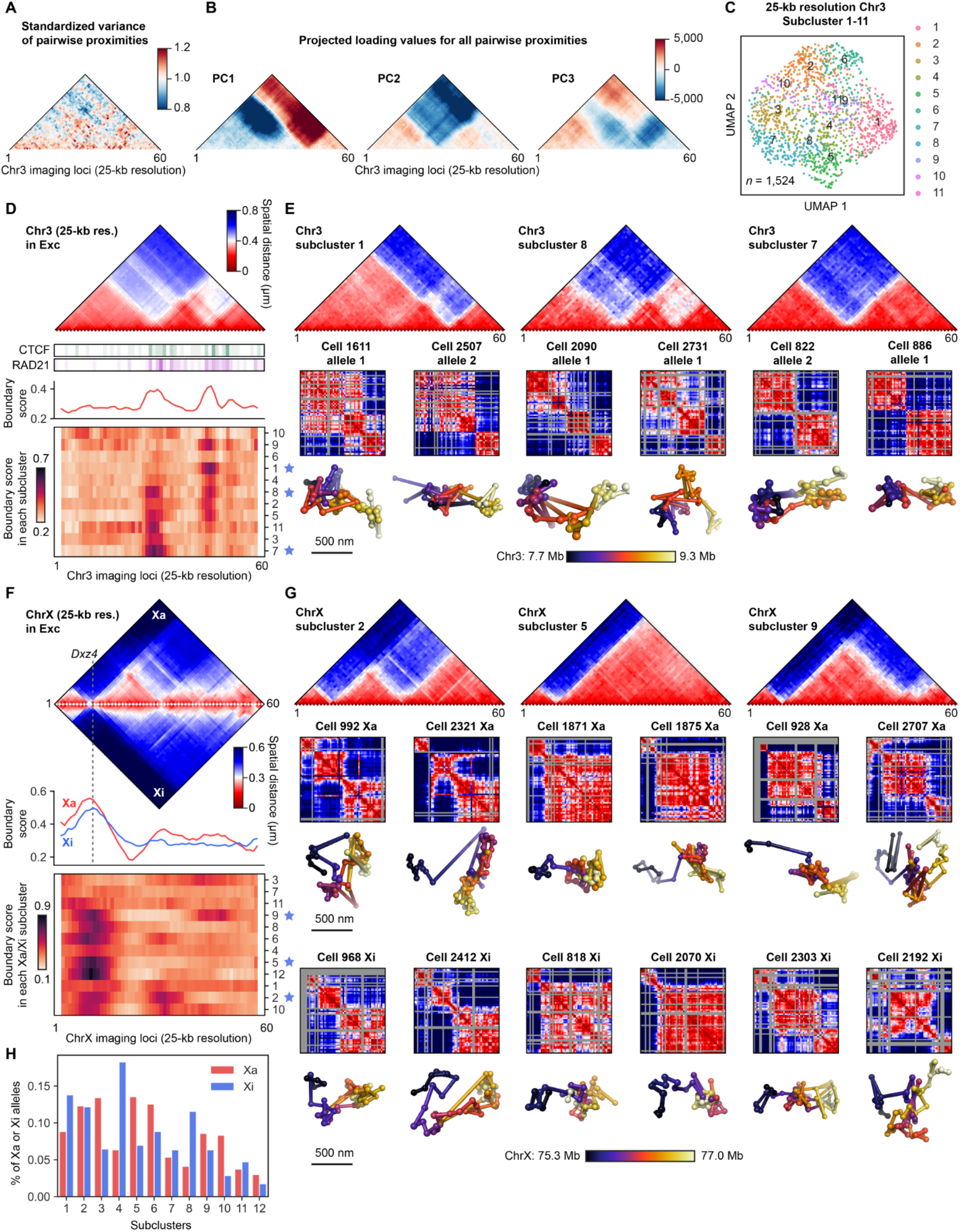
Heterogeneous domain structures in single chromosomes. (**A** and **B**) Principal component analysis (PCA) on the chromosome 3 region (7.7-9.3 Mb) that was targeted at 25-kb resolution. The variance of the pairwise spatial proximity map is shown in (A) with the corresponding PCA loading shown in (B). (**C**) UMAP visualization of individual chromosomes in 11 structural subclusters for the 25-kb resolution chromosome 3 data (n = 1,524 chromosomes) from excitatory neurons. (**D**) Bulk averaged pairwise distance map from excitatory neurons (top) shows distinct domains in the chromosome 3 region with boundaries and high median boundary score regions matching CTCF and cohesin binding sites obtained by high-throughput sequencing (ChIP-seq) with the mouse brain (*77*). Single chromosomes were clustered into different domain structures with distinct median boundary score profiles. (**E**) Examples for three of the structural subclusters are shown with different pairwise distance map structures. Single chromosome distance maps and chromosome structures from individual cells are shown below for each subcluster. Chromosomes with >20% loci detected in n = 1,895 excitatory neurons from three biological replicates were used in (A)-(E). (**F**) Pairwise median distance maps for Xa (top, upper triangle) and Xi (top, lower triangle) in excitatory neurons from the 25-kb DNA seqFISH+ data. Clustering of the domain structures for individual Xa and Xi shown with the median boundary scores for Xa and Xi (middle) and for each subcluster containing Xa and Xi (bottom) in excitatory neurons from the 25-kb DNA seqFISH+ data. The location for the macrosatellite DXZ4 locus at the boundary of two mega-domains in Xi (*83*) is shown (top and middle). (**G**) Examples for subclusters 2, 5 and 9 with corresponding single-cell pairwise distance maps and single-cell 3D chromosome structures for Xa and Xi in excitatory neurons. Individual Xa and Xi can have similar domain structures. (**H**) The frequency of each subcluster occurring in single cells for Xa and Xi from 805 excitatory neurons. n = 805 cells with >20% loci detected for both Xa and Xi regions (75.3-77.0 Mb) from three biological replicates in (F)-(H).

These results demonstrate that even in cells expressing CTCF and cohesin, there are highly variable domain boundaries and inter- and intra-domain interactions that can be obscured with ensemble-averaged measurements. Recent super-resolution imaging studies (*33, 34*) have shown that CTCF and cohesin have differential effects on chromosomal domain formation in single cells, where CTCF preferentially promotes intra-domain interactions whereas cohesin promotes stochastic intermingling of inter-domains. In addition, cohesin appears to alter the instantaneous transcription activities of boundary-proximal genes (*33*). It is possible that the variabilities of single-cell domains we observed (Fig. 6 and fig. S6 and S7) are mediated by stochastic combinatorial binding of the architectural proteins (*78*) such as CTCF and cohesin at individual chromosomes and may lead to instantaneous transcriptional differences at those domain boundaries in single cells.

### Active and inactive X chromosome organization in single cells

Lastly, we examined the differences of chromatin states and chromosome conformations between the active X chromosome (Xa) and the inactive X chromosome (Xi) from the female mouse brain cortex. X-chromosome inactivation in female mammalian cells has been extensively studied as a model system for chromosome organization (*59*). The imaging-based genomics data can straightforwardly distinguish the Xa and Xi based on their mutually exclusive associations with Xist RNA, a long noncoding RNA that is specifically expressed from and associates with the Xi (*59*) (fig. S8A and S8B). As expected, the Xi showed enrichment of repressive mH2A1 and H3K27me3 marks (*79, 80*), high nucleolar association (*80*), depletion of active chromatin mark (*81*) H3K4me2, and condensed LINE1 DNA elements (*82*) (fig. S8A and S8B).

To gain insight into the structural differences between the Xa and Xi, we calculated the median spatial distance and spatial proximity maps for the whole X chromosome at 1-Mb resolution as well as for a targeted region (75.3-77.0 Mb) at 25-kb resolution in a cell-type-specific fashion (fig. S8C to F). We observed that the Xa and Xi have distinct median distances between pairs of loci at different genomics length scales and in different cell types (fig. S8G and S8H). In particular, we observed that the Xi is organized into two mega-domains separated at the macrosatellite DXZ4 locus at the whole chromosome scale, consistent with the literature (*83*). Interestingly, although the Xi is more compact than the Xa globally at the larger scale of tens of megabases (fig. S8C and S8G) (*22, 84*), we found the Xa appears to be more structured and compact at the Mb or below length scales based on the population-averaged spatial distance quantification (fig. S8C, D, G and H).

To further resolve the observed bulk structural differences between the Xa and Xi, we examined the Xa and Xi conformation systematically at the single-cell level by applying the PCA-based approach used for the autosomal regions (Fig. 6A to E and fig. S6 and S7). Interestingly, both the Xa and Xi have heterogeneous domain structures in individual cells (Fig. 6F and 6G and fig. S9). Even the region of the Xi that appears unstructured in the bulk data from both DNA seqFISH+ (Fig. 6F and fig. S8D) and allele-specific Hi-C studies (*83, 85*) appeared to adopt discrete domains in subsets of cells (Fig. 6G). We found that similar domain subclusters can be used in both the Xa and Xi, but with different relative frequencies for specific chromosome conformation (Fig. 6H), which leads to different average conformations for the Xa and Xi (Fig. 6F and fig. S8D). Taken together, although the Xa and Xi show very different chromatin states and ensemble-averaged chromosome conformation, they can share similar underlying single-cell domain structures with different relative conformational preferences at the finer scale, which has been obscured with ensemble-averaged bulk measurements.

## Discussion

Our work demonstrates cell type-dependent and -independent features in nuclear organization across thousands of single cells in the mouse cerebral cortex, derived by integrated spatial genomics tools to measure RNA, DNA and chromatin marks. In particular, we examined nuclear morphologies, global chromatin states, DNA-nuclear body associations, radial nuclear organization, as well as chromosome and domain structures in transcriptionally defined cell types. Existing microscopy and sequencing-based datasets showed high concordance with our results at each modality, supporting the robustness and accuracy of our approach and allowing the exploration of multimodal nuclear features in tissue sections.

At the global level, our observations indicate that the chromosome organization in a cell reflects cell-type specific nuclear body arrangements. We observed that each chromosome contains a unique pattern of fixed points associated with each nuclear body and chromatin mark such that the DNA loci are reliably located on the exterior of the nuclear bodies in single cells at the scale of 1 Mb. Some of the fixed points are cell type-dependent, whereas others, such as nuclear speckle-associated loci (*68*), are largely cell type-independent. Because nuclear body morphologies are different in different cell types, these fixed points lead to distinct organizations of the nucleus in each cell type. For example, because most Pvalb inhibitory neurons have a large central heterochromatic globule, chromosomes 7 and 17 with fixed points to heterochromatin are organized around this central hub. These chromosomes are often found in the nuclear interior and interact with many other chromosomes. We had observed both nuclear bodies and fixed points in mouse ES cells (*32*), and now show that this principle operates in tissues to drive cell type-specific genome organization. Similar observations of radial chromosome and nuclear body reorganization during brain and retinal development (*18, 19, 61, 74*) suggest the same principle could also be applied to the developmental processes.

At the 25-kb scale organization of chromosomes, the single-cell resolution of the DNA seqFISH+ data explains that nested structures and regions with ambiguous boundaries that are often observed in bulk Hi-C contact maps (*86*) are due to different domains appearing in individual cells, possibly because subsets of CTCF and cohesin sites are stochastically used to insulate individual chromosomes. In line with this observation, single-cell domain structures were previously observed even in cohesin depleted cells where domain structures were abolished at the population-averaged level, suggesting the population level domains are formed due to preferential domain boundary positions (*23*). Our systematic single-cell analysis further showed the prevalence of clear single-cell domains even in regions which lack ensemble domain boundaries such as the Xi region (*83, 85*), demonstrating the importance of studying chromosome structures in individual cells to better interpret the organizational principles of 3D genome architecture.

The robust demonstration of integrated spatial genomics in tissues indicates the same approach can be applied to a diverse range of biological systems to further explore the diversity and invariant nuclear architectures in single cells. At the global scale of nuclear organization, it is still unclear how the distinct nuclear body and chromosome arrangements as well as their associations arise in the first place in different cell types. In addition, given the prevalence of the diverse single-cell domain structures that can differ from ensemble-averaged TADs, it would be critical to study both single-cell domain structures and instantaneous transcriptional activity in each domain from the same single cells and understand their relationships at a fine spatiotemporal resolution. We anticipate addressing these questions in future studies by genome-scale chromosome imaging together with transcriptome-scale profiling (*53, 87*) and protein imaging as well as by “track first and identify later” live-cell approaches (*88, 89*) with multiplexed chromosome labeling (*90, 91*).

## Acknowledgements

We thank N. Ollikainen and M. Guttman for help with Hi-C analysis; R.J. Oakey for providing the processed ChIP-seq data; C.-H.L. Eng for helpful discussion; C. Karp for making custom-made flow cells; L. Sanchez and E. Buitrago-Delgado for help with the perfusion setup; I. Strazhnik for help with figures; A. Anderson for help with the manuscript.

## Author contributions

Y.T. and L.C. conceived the idea and designed experiments. Y.T. designed probes with input from C.T.. Y.T., J.Y., and S.Schindler prepared and validated experimental materials. Y.T. performed imaging experiments. Y.T., S.Shah., N.P., and J.W. wrote image processing scripts, and Y.T. performed image processing. Y.T., S.Z., and L.C. analysed data with input from G.-C.Y.. Y.T. and L.C. wrote the manuscript with input from S.Z., C.T., and G.-C.Y.. L.C. supervised all aspects of the projects.

## Data and materials availability

The custom written scripts used in this study are available in a Github repository (https://github.com/CaiGroup/dna-seqfish-plus-tissue). The source data and processed data from this study are available in a Zenodo repository (*55*).

## Materials and Methods

### Primary and readout probe design and synthesis

RNA seqFISH primary probes for marker genes were designed similarly to our previous studies(*32, 53, 87*). In brief, individual RNA species were encoded in fluorescent channel 1 (635 nm) and 2 (561 nm) at each hybridization round and sequentially called. To implement this, 35-nt RNA target binding sequences, four 15-nt unique readout probe binding sites encoded for each RNA target, and a pair of 20-nt primer binding sites at the 5ʹ and 3ʹ ends of the probe for probe generation were concatenated. Marker genes were selected based on previous studies with mouse brain samples(*46–50, 52, 53, 92*).

DNA seqFISH+ encoding strategy and primary probe design with GRCm38/mm10 mouse genome are described in detail in our previous study(*32*). In brief, a total of 80 serial rounds as a 16-base coding scheme with 5 rounds of barcoding was encoded in fluorescent channel 1 (635 nm) and 2 (561 nm) for the 1-Mb resolution data using a subset of barcodes in the codebook (table S1). In addition, a combined strategy of diffraction limited spot imaging (60 serial rounds) and chromosome painting (20 serial rounds) was encoded in fluorescent channel 3 (488 nm) to resolve 20 distinct regions (1.5-2.4-Mb in size) with 25-kb resolution. In total, 3,660 loci were targeted including 2,460 loci at 1-Mb resolution for 20 chromosomes and 1,200 loci at 25-kb resolution for the specific 20 distinct regions. To implement this, 35-nt genomic DNA target binding sequences, five 15-nt unique readout probe binding sites encoded for each barcoding round, and a pair of 20-nt primer binding sites at 5ʹ and 3ʹ end of the probe for probe generation were concatenated.

The repetitive element DNA FISH probes (LINE1, SINEB1, Telomere, MinSat) were designed as described before(*32*). Similarly to Telomere and MinSat probes, the MajSat probe (Integrated DNA Technologies) was designed as a dye-conjugated 15-nt probe using the following sequence (5ʹ-TGTCCACTGTAGGAC), which directly targeted genomic DNA. In addition, rDNA primary probes (a total of 101 probes) targeting 45S pre rRNA gene (GenBank: X82564.1) were designed similarly to DNA seqFISH+ probes except for containing four of 15-nt unique readout probe binding sites specific to rDNA primary probes.

RNA seqFISH and DNA seqFISH+ primary probes were generated from oligoarray pools (Twist Bioscience) with enzymatic amplifications as previously described(*32*) based on Oligopaint technologies(*21*).

RNA seqFISH, DNA seqFISH+ and sequential immunofluorescence readout probes (5’ amine-modified DNA oligonucleotides, Integrated DNA Technologies) of 12-15-nt in length were designed and conjugated to Alexa Fluor 647-NHS ester (Invitrogen A20006) or Cy3B-NHS ester (GE Healthcare PA63101) or Alexa Fluor 488-NHS ester (Invitrogen A20000) in-house as described previously(*32*).

### DNA-antibody conjugation

Preparation of the oligonucleotide-conjugated antibodies was performed, as described previously(*32, 58*). The BSA-free primary antibodies were purchased from commercial vendors as listed below. As a conjugation strategy, the crosslinking of 5’ thiol-modified 18-nt DNA oligonucleotides (Integrated DNA Technologies) to lysine residues on antibodies was performed via PEGylated SMCC cross-linker (SM(PEG)2) (Thermo Scientific Thermo Scientific 22102). As an alternative conjugation strategy, SiteClick R-PE Antibody Labeling Kit (Life Technologies S10467) was also used to crosslink 5’ DBCO-modified 18-nt DNA oligonucleotides (Integrated DNA Technologies) to the specific sites on primary antibodies. The oligonucleotide-conjugated primary antibodies were individually validated by SDS-PAGE gel and immunofluorescence, and stored in 1× PBS at −80 °C as small aliquots.

The oligonucleotide DNA-conjugated primary antibodies used were as follows: mH2A1 (Abcam ab232602), H3K27ac (Active Motif 39133), H3K27me2 (Cell Signaling 9728BF), H3K27me3 (Cell Signaling 9733BF), H3K4me2 (Cell Signaling 9725BF), H3K9me3 (Diagenode MAb-146-050), H4K20me3 (Active Motif 39671), MeCP2 (Cell Signaling 3456BF), RNA polymerase II (phospho S5) (Abcam ab5408), SF3a66 (Abcam ab77800). Two antibodies (H3K27ac, RNA polymerase II (phospho S5)) were excluded from the downstream analysis owing to the signal dimness (H3K27ac) and incomplete penetration of the antibody (RNA polymerase II (phospho S5)).

### Tissue slice preparation

All animal care and experiments were carried out in accordance with Caltech Institutional Animal Care and Use Committee (IACUC) and NIH guidelines. 6-7-week-old C57BL/6J female mice were obtained from The Jackson Laboratory.

To attach tissue sections, the 24 × 60 mm coverslips (VWR 16004-312) were cleaned by sonication in 1 M sodium hydroxide and 100% ethanol in Ultrasonic Cleaner (Fisher Scientific FS20) three times for 10 minutes each, followed by a 15 minute sonication in 100% acetone. The coverslips were then immersed in 2% (v/v) (3-Aminopropyl)triethoxysilane (Sigma A3648) prepared in acetone for 2 minutes at room temperature. Then the coverslips were rinsed twice in water and heat treated at 90°C for 20 minutes. Next the coverslips were treated with 90 µg/ml of Poly-D-lysine (Sigma P7280) in 1× PBS (Invitrogen AM9625) for 16 hours at room temperature, followed by 3 times rinsing in nuclease-free water. The coverslips were then air-dried and attached to a microscope slide (VWR 48312-004) to facilitate the collection of cryo-sections. The coverslips were freshly prepared as required for sectioning below.

Mice were perfused for 4 minutes with perfusion buffer (10 U/ml Heparin (Sigma-Aldrich H3149), 0.5% Sodium nitrite (w/v) (Sigma-Aldrich 237213) in 1xPBS) upon isoflurane anesthesia, followed by fresh 4% PFA (EMS 15714) in 1× PBS buffer for 4 minutes with a flow rate of 5 mL/min through the peristaltic pump (Masterflex MP-07557-00). The mouse brains were dissected out of the skull and immediately placed in 4% PFA in 1× PBS for 16 hours at 4°C. The brains were then immersed in 10% (w/v) RNAse-free Sucrose (Amresco 0335–2.5KG) in 1x PBS for 1 hour at room temperature, then 20% (w/v) RNAse-free Sucrose in 1xPBS, until the brains sank. Next the samples were incubated for 16 hours in 30% (w/v) RNase-free Sucrose in 1× PBS at 4°C. After the brains sank, they were embedded in OCT (Sakura 4583) individually and frozen in a bath of dry ice and ethanol. The samples were stored at −80°C until 10-15 μm coronal sections were cut using a cryostat (Leica CM3050). The sections were immediately placed on the functionalized coverslips described above. The tissue slices were stored at −80°C until the tissue slice experiment.

### Tissue slice experiment

The combined sequential immunofluorescence, RNA seqFISH and DNA seqFISH+ sample preparation was performed similarly to our previous study(*32*) with some modifications for tissue slice experiments. The tissue slice samples were dried, and permeabilized with 70% ethanol pre-chilled to −20°C at room temperature for 15 minutes. The samples were then dried and further permeabilized with 8% Triton-X (Sigma-Aldrich 93443) in 1× PBS at room temperature for 30 minutes after attaching a sterilized silicon plate (McMASTER-CARR 86915K16) with a punched hole to the coverslip to use it as a chamber. The samples were washed three times with 1× PBS and blocked at room temperature for 15 minutes with blocking solution consisting of 1× PBS, 10 mg/mL UltraPure BSA (Invitrogen AM2616), 0.3% Triton-X, 0.1% dextran sulfate (Sigma D4911) and 0.5 mg/mL sheared Salmon Sperm DNA (Invitrogen AM9680). Then oligonucleotide DNA-conjugated primary antibodies (see ‘DNA-antibody conjugation’) with an estimated each concentration of 1-5 ng/μl were incubated in the blocking solution with 100-fold diluted SUPERase In RNase Inhibitor (Invitrogen AM2694) at room temperature for 18-24 hours. The samples were washed with 1× PBS three times and incubated at room temperature for 15 minutes, fixed with freshly made 4% formaldehyde in 1× PBS at room temperature for 5 minutes, and washed with 1× PBS six times and incubated at room temperature for 15 minutes. The samples were then further fixed with 1.5 mM BS(PEG)5 (PEGylated bis(sulfosuccinimidyl)suberate) (Thermo Scientific A35396) in 1× PBS at room temperature for 30 minutes, followed by quenching with 100 mM Tris-HCl pH7.4 (Alfa Aesar J62848) at room temperature for 5 minutes. Then the samples were washed with 1xPBS and 70% ethanol three times, and air dried by removing the custom silicon chamber.

After the immunofluorescence preparation steps, custom-made flow cells (fluidic volume about 40 μl), which were made from glass slide (25 × 75 mm) with 1-mm thickness and 1-mm diameter holes and a PET film coated on both sides with an acrylic adhesive with total thickness 0.25 mm (Grace Bio-Labs RD481902), were attached to the coverslips. The samples were rinsed three times with a 50% denaturation buffer consisting of 50% formamide (Invitrogen AM9342) and 2× SSC, and incubated at room temperature for 15 minutes. Then the samples were optionally heated at 80°C for 3 minutes (this step was performed for replicate 1 and 2, and omitted for replicate 3), and washed with 2× SSC twice. Then RNA seqFISH primary probe pools (1-10 nM per probe) including mRNA, intron and non-coding RNA targets were hybridized in a 50% hybridization buffer consisting of 50% formamide, 2× SSC and 10% (w/v) dextran sulfate (Millipore 3710-OP). The hybridization was performed at 37°C for 3 days in a humid chamber. After the hybridization step, the samples were washed with a 40% wash buffer consisting of 40% formamide, 2× SSC and 0.1% Triton X-100 at 37°C for 15 minutes, followed by three rinses with 4× SSC, and stored at 4°C until imaging. Then the samples were imaged as described below (see ‘seqFISH imaging’). Note that H4K20me3 immunofluorescence imaging was also performed at this step for validation and alignment.

After the completion of RNA seqFISH imaging, the samples were prepared for DNA seqFISH+ imaging. The samples were rinsed with 1× PBS, and incubated with 100-fold diluted RNase A/T1 Mix (Thermo Fisher EN0551) in 1× PBS at 37°C for 1 hour. Then samples were rinsed three times with 1× PBS, followed by three rinses with the 50% denaturation buffer and an incubation at room temperature for 15 minutes. Then the samples were heated on the heat block at 90°C for 7 minutes in the 50% denaturation buffer, by sealing the holes of the custom chamber with aluminum sealing tapes (Thermo Scientific 232698). After the heating, the samples were rinsed with 2× SSC and hybridized with a powder of DNA seqFISH+ primary probe pools, which were dried by speed-vac and consisted of about 1 nM each DNA seqFISH+ probe, 1 μM LINE1, 1 μM SINEB1 and 100 nM 3632454L22Rik locus fiducial marker probes(*32*), resuspended in a 40% hybridization buffer consisting of 40% formamide, 2× SSC and 10% (w/v) dextran sulfate. For the rDNA FISH experiment, about 20 nM each rDNA probe and 100 nM 3632454L22Rik locus fiducial marker probe were resuspended in the 40% hybridization buffer. The DNA seqFISH+ primary probe hybridization was performed at 37°C for 3-5 days in a humid chamber. After hybridization, the samples were washed with a 30% wash buffer consisting of 30% formamide, 2× SSC and 0.1% Triton X-100 at room temperature for 15 minutes, followed by three rinses with 4× SSC.

Then samples were further processed to ‘padlock’ DNA seqFISH+ primary probes on the genomic DNA in order to increase the stability of primary probes and prevent the loss of signals during 80 rounds of DNA seqFISH+ imaging routines (see ‘seqFISH imaging’). A 31-nt global ligation bridge oligonucleotide (Integrated DNA Technologies, 5ʹ-TCAGTTGCAGCGCATGCTCGACCAAGGCTGG) was hybridized in a 20% hybridization buffer consisting of 20% formamide, 10% dextran sulfate (Sigma D4911) and 4× SSC at 37°C for 2 hours. The global ligation bridge was designed to hybridize to the 15-nt sequence of the DNA seqFISH+ primary probes at the 5ʹ end and the 16-nt sequence at the 3ʹ end. Then, samples were washed with a 12.5% wash buffer consisting of 12.5% formamide, 2× SSC and 0.1% Triton X-100 three times and incubated at room temperature for 5 minutes, followed by three rinses with 1× PBS. The samples were then incubated with 20-fold diluted Quick Ligase in 1× Quick Ligase Reaction Buffer from Quick Ligation Kit (NEB M2200) supplemented with an additional 1 mM ATP (NEB P0756) at room temperature for 1 hour to allow ligation reaction between the 5ʹ- and 3ʹ-ends of the DNA seqFISH+ primary probes. Then the samples were washed with the 12.5% wash buffer, followed by three rinses with 1× PBS.

The samples were further processed for amine modification and post-fixation to increase the stability of the primary probes across imaging rounds along with the ligation(*32*). First, the samples were rinsed with 1× labelling buffer A, followed by incubation with tenfold diluted Label IT amine modifying reagent in 1× labelling buffer A from Label IT nucleic acid modifying reagent (Mirus Bio MIR 3900) at room temperature for 45 minutes. Second, after three rinses with 1× PBS, the samples were fixed with 1.5 mM BS(PEG)5 in 1× PBS at room temperature for 30 minutes, followed by quenching with 100 mM Tris-HCl pH7.4 at room temperature for 5 minutes. Then the samples were washed with the 55% wash buffer at room temperature for 5-15 minutes, followed by three rinses with 4× SSC, and stored at 4°C until imaging. Then the samples were imaged for DNA seqFISH+ and sequential immunofluorescence as described below (see ‘seqFISH imaging’).

After the completion of DNA seqFISH+ and sequential immunofluorescence imaging, Concanavalin A (ConA) staining and imaging(*63*) were performed for nuclear segmentation. The samples were incubated with a ConA solution consisting of 20-100 μg/mL Alexa Fluor 488 conjugate of ConA (Invitrogen C11252), 1× PBS and 0.3% Triton X-100 at room temperature for 2 hours, and washed with the 55% wash buffer at room temperature for 2 minutes, followed by a rinse with 4× SSC. Then the samples were imaged as described below (see ‘seqFISH imaging’).

### Cell culture experiment

E14 mouse ES cells (E14Tg2a.4) from Mutant Mouse Regional Resource Centers were maintained under serum/LIF condition as previously described(*32, 87*). NIH/3T3 cells (ATCC CRL-1658) were cultured as previously described(*53, 93*).

The combined sequential immunofluorescence, RNA seqFISH and DNA seqFISH+ protocol was performed as previously described(*32*) for E14 mouse embryonic stem cells and NIH/3T3 mouse fibroblast cell line with ITS1 RNA FISH probe and rDNA FISH probes used in the tissue slice experiments.

### Image acquisition

#### Microscope setup

All imaging experiments were performed with the confocal fluorescence imaging platform and fluidics delivery system similar to previous studies(*32, 53, 87*). The microscope (Leica DMi8) was equipped with a confocal scanner unit (Yokogawa CSU-W1), a sCMOS camera (Andor Zyla 4.2 Plus), a 63× oil objective (NA = 1.40, Leica 11506349), and a motorized stage (ASI MS2000). Fiber coupled lasers (635, 561, 488 and 405 nm) from CNI and Shanghai Dream Lasers Technology and filter sets from Semrock were used. The custom-made automated sampler was used to move to the well of the designated hybridization buffer corresponding to each hybridization round from a 2.0-mL 96-well plate (Corning 3960) and hybridization buffers were moved through a multichannel fluidic valve (IDEX Health & Science EZ1213-820-4) to the custom-made flow cell using a syringe pump (Hamilton Company 63133-01). Other buffers used for the imaging routine were also moved through the multichannel fluidic valve to the custom-made flow cell using the syringe pump. The control of imaging and the automated fluidics delivery system was achieved by a custom-written script in μManager(*94*).

#### seqFISH imaging

The sequential hybridization and imaging routines were performed similarly to those previously described(*32, 53, 87*) with some modifications. Briefly, the sample with the custom-made flow cell was first connected to the automated fluidics system on the motorized stage on the microscope. Fields of view (FOVs) (4 FOVs in replicate 1, 5 FOVs in replicate 2, and 8 FOVs in replicate 3 of the mouse cortex) were registered using nuclei signals stained with DAPI solution consisting of 5 μg/mL DAPI (Sigma D8417) and 4× SSC. First, RNA seqFISH and H4K20me3 immunofluorescence imaging was performed with the sequential hybridization and imaging routines described below. After the completion of the imaging, the samples were disconnected from the microscope, and proceeded to the DNA seqFISH+ procedures (see ‘Tissue slice experiment’). For the DNA seqFISH+ and sequential immunofluorescence imaging, the registered FOVs for RNA seqFISH were loaded and manually shifted to image the same FOVs as RNA seqFISH imaging. The DNA seqFISH+ and sequential immunofluorescence imaging was performed with the sequential hybridization and imaging routines described below, followed by ConA staining and imaging.

All the following sequential hybridization and imaging routines were performed at room temperature on the microscope. For the sequential hybridization routine, the serial hybridization buffer consisting of two or three unique readout probes (10-50 nM) with different fluorophores (Alexa Fluor 647, Cy3B or Alexa Fluor 488) and 10% EC buffer (10% ethylene carbonate (Sigma E26258), 10% dextran sulfate (Sigma D4911) and 4× SSC) was picked up from a 96-well plate and incubated with the sample for 20 minutes. After the serial hybridization buffer incubation, the samples were washed with 1 mL of a 4× SSCT buffer consisting of 4× SSC and 0.1% Triton-X, followed by a wash with 330 μL of the 12.5% wash buffer. Then, the samples were rinsed with about 200 μL of 4× SSC, followed by a staining with about 200 μL of the DAPI solution for 30 seconds. Next, an anti-bleaching buffer consisting of 50 mM Tris-HCl pH 8.0 (Invitrogen 15568025), 4× SSC, 3 mM Trolox (Sigma 238813), 10% D-glucose (Sigma G7528), 100-fold diluted catalase (Sigma C3155), 1 mg/mL glucose oxidase (Sigma G2133) was flowed through the sample for imaging. The anti-bleaching buffer was covered by a mineral oil (Sigma-Aldrich M5904) to prevent the progress of the enzymatic reaction in the tube. After image acquisition described below, 1 mL of the 55% wash buffer was flowed for 1 minute to strip off readout probes, followed by an incubation for 1 minute and rinsing with 4× SSC. The above serial hybridization, imaging and signal extinguishing steps were repeated until the completion of all rounds. For the RNA seqFISH and DNA seqFISH+ experiments, blank images containing only autofluorescence of the tissue section were imaged at the beginning and end of the routines. For the DNA seqFISH+ experiments, fiducial marker images containing only fiducial markers were obtained at the beginning and end of the routines for image registration.

For the imaging routine, snapshots were acquired per fluorescent channel per field of view with 0.75 μm z-steps over 10 μm z-slices for RNA seqFISH for mRNA and intron targets, and with 0.25 μm z-steps for RNA seqFISH for non-coding RNA targets (ITS1, Malat1, Xist), DNA seqFISH+, sequential immunofluorescence and ConA staining. RNA seqFISH imaging was performed with 635 nm, 561 nm, 488 nm fluorescent channels except for a DAPI alignment hybridization round in the end with 635 nm, 561 nm, 488 nm and 405 nm fluorescent channels. During RNA seqFISH, 635 nm and 561 nm fluorescent channels contained RNA seqFISH targets while a 488 nm fluorescent channel contained polyA staining for an alignment. Importantly, we omitted the DAPI imaging during RNA seqFISH imaging routines to prevent the nuclear damage with 405 nm laser exposure prior to DNA seqFISH+ preparation in the tissue sections. DNA seqFISH+ imaging was performed with 635 nm, 561 nm, 488 nm and 405 nm fluorescent channels, containing DNA seqFISH+ targets in the first 3 fluorescent channels and DAPI staining in the last 405 nm fluorescent channel. The fiducial markers were also included in the 3 fluorescent channels to allow image registration at the subpixel resolution. Sequential immunofluorescence imaging was performed with 635 nm, 561 nm, 488 nm and 405 nm fluorescent channels, containing primary antibody targets in the first 2 fluorescent channels and DAPI staining in the last 405 nm fluorescent channel. At the end of all imaging routines, images were manually checked to repeat problematic hybridization rounds such as off-focus and intensity saturation. In total, it took approximately 3-4 days to complete the 80 rounds of the hybridization and imaging routine under our DNA seqFISH+ conditions.

### Image analysis

The image analysis was performed using ImageJ (v1.51s), Python (v3.7.4), MATLAB R2019a and ilastik (v1.3.3)(*95*) as described previously(*32*) with modifications and implementation of 3D nuclear segmentation.

#### 3D nuclear segmentation

DAPI images from the first round of DNA seqFISH+ imaging and ConA images from the last round of imaging were aligned and chromatic shifts were corrected. The images were preprocessed for training by binning the image down by a factor of 4 in the x and y dimension with the sum intensity. The preprocessed images were imported into the pixel classification module in ilastik(*95*), an interactive supervised machine learning software, and trained using all possible features. For the DAPI images, the interior, edge and exterior of the DAPI signal were used as classifiers. For the ConA images, the ConA signal, ConA signal edge, and ConA signal holes (location of nuclei) were trained as classifiers. The probability maps for each position were output as hdf5 files, which were then read into MATLAB. The nuclear edge probability was defined as the probability of ConA edge times the probability of DAPI signal edge time 1 minus the probability of DAPI interior time 1 minus the probability of ConA exterior. Seeds for the seeded watershed algorithm were created by multiplying the DAPI interior probability with 1 minus the probability of ConA edge and setting an appropriate probability threshold. Finally, background pixels were identified as DAPI exterior probability above an appropriate threshold. The array for watershed was then formed by taking the edge probability array and creating global minima at the seed and background pixels. A watershed algorithm was then performed on the resulting array. Objects smaller than an appropriate area and objects greater than an appropriate area were removed from the resulting segmentation to achieve a final nuclear segmentation.

#### Preprocessing of images

A flat field correction was applied by dividing the normalized background illumination with each of the fluorescence images to correct the non-uniform background intensities while preserving the intensity profile of the fluorescent points whenever necessary. The background signal was subtracted using the ImageJ rolling ball background subtraction algorithm with a radius of 3 pixels for RNA seqFISH (mRNA and intron targets) and DNA seqFISH+ images. The background subtraction was not performed for non-coding RNA and immunofluorescence images except for the foci detection analysis described below.

#### FISH spot detection and fitting

RNA seqFISH and DNA seqFISH+ spot locations were obtained as described previously(*32*) by using a Laplacian of Gaussians filter, semi-manual thresholding and a 3D local maxima finder. Subsequently the DNA seqFISH+ spot locations were super resolved using a 3D radial center algorithm(*96, 97*) found on the Parthasarathy lab website (https://pages.uoregon.edu/raghu/particle_tracking.html).

#### Correction of chromatic effects and alignment

Chromatic aberration shifts between different fluorescent channels were corrected by applying the calculated offsets using the fiducial markers or 0.2 μm TetraSpeck Microspheres (Thermo Scientific T7280) appeared in each fluorescent channel.

All images were aligned to the initial hybridization (hyb1) image in DNA seqFISH+. To align RNA seqFISH images in different hybridization rounds, reference channels (polyA staining in 488 nm fluorescent channel) were first aligned to RNA seqFISH hyb1 image using 3D phase correlations along every axis iteratively to find a consensus transformation for alignment as previously described(*87*). The aligned DAPI image taken at the last round of RNA seqFISH imaging was further aligned to DNA seqFISH+ hyb1 image and the obtained transformation was propagated to all the other RNA seqFISH images, considering the differences of z-sampling (0.75 μm z-steps for RNA seqFISH and 0.25 μm z-steps for DNA seqFISH+). Then further alignment to correct any rotation between RNA seqFISH and DNA seqFISH+ images was done as previously described(*32*) using DAPI and H4K20me3 staining taken at both RNA seqFISH and DNA seqFISH+ imaging routines. Similarly, to align sequential immunofluorescence images in different hybridization rounds, reference channels (DAPI staining in 405 nm fluorescent channel) were aligned to DNA seqFISH+ hyb1 image using 3D phase correlations. To align DNA seqFISH+ spots in different hybridization rounds at subpixel resolution, identified fiducial markers in each fluorescent channel (635 nm, 561 nm and 488 nm) were used in Python as described previously(*32*) with modified parameters.

#### Decoding for RNA seqFISH

The identified spots within individual 3D nuclear ROIs obtained by the 3D nuclear segmentation step were collected, and RNA identities were resolved by checking the identity of the hybridization round and fluorescent channel.

#### Decoding for DNA seqFISH+

Both 1-Mb resolution (635 nm and 561 nm fluorescent channels) and 25-kb resolution (488 nm fluorescent channel) DNA seqFISH+ decoding were performed as described previously(*32*). Decoding was performed at each entire field of view, and then the identified spots within individual 3D nuclear ROIs were collected.

#### Nuclear marker image analysis

Raw intensity values for all the voxels within individual 3D nuclear ROIs were obtained for all immunofluorescence raw images as well as repetitive element DNA (LINE1, SINEB1, Telomere, MajSat, MinSat and rDNA), non-coding RNA (ITS1, Malat1 and Xist) and DAPI raw images, and exported as csv files.

The edge detection for chromatin marker exterior quantification was performed as described previously(*32*). In brief, Find edges function in ImageJ was performed on background subtracted images (rolling ball radius 3 pixels), and then the intensity values were obtained similarly to the raw images.

The foci detection for chromatin markers was performed with Yen’s auto threshold method in ImageJ on background subtracted images (rolling ball radius 9 pixels) for each slice in the z-stack to binarize the images, followed by filling and opening of the binary images to remove internal voids and shot noise. After objects smaller than a voxel of 20 or greater than a voxel of 100,000 were removed, the remaining objects were labeled with unique numbers, which correspond to individual nuclear marker foci. The labeled foci within individual 3D nuclear ROIs were exported as csv files.

#### Conversion of voxel information to physical distance

After image analysis steps above, voxel information was converted to physical distance based on our microscope setup and imaging condition with 0.103 μm for x and y and 0.250 μm (or 0.750 μm in mRNA and intron seqFISH) for z voxels for the subsequent downstream data analysis.

### Hi-C data analysis

Hi-C data for mouse cortex(*6, 56*) was obtained from NCBI GEO (accession GSE35156) and was processed using Juicer tools(*98*). Contact maps containing Knight–Ruiz normalized counts(*99*) were obtained. Hi-C data were binned at 25-kb and 1-Mb resolution, and overlapping regions within a given bin size for 1-Mb resolution were excluded from the analysis. Note that while the Hi-C data was originally used with 40-kb binning(*6, 56*), we used it with higher resolution of 25-kb binning to directly compare to our 25-kb resolution DNA seqFISH+ data.

### CTCF and Rad21 ChIP-seq data analysis

Processed CTCF and Rad21 ChIP-seq data in mouse brain(*77*) were kindly provided by Rebecca J. Oakey. Peaks were identified using peak shifts and window sizes of 138 bp and 144 bp for CTCF and Rad21, respectively with a False Discovery Rate below 10%(*77*). The mm10 genomic coordinates for the DNA seqFISH+ loci were converted to mm9 using the UCSC Genome Browser LiftOver tool and were compared to the ChIP-seq peaks.

### RNA seqFISH data analysis

The mRNA counts in each nuclear segmentation were normalized within each cell by the total Eef2 mRNA counts. Then each gene is z-score normalized across all the cells. Hierarchical clustering was then performed on the normalized cell by gene matrix using Mathematica function “Agglomerate” with Ward distance. The clustering results obtained above were visualized with uniform manifold approximation and projection (UMAP)(*100*) using a UMAP-learn library in Python. The cell type annotation is based on the top differentially expressed genes. Two excitatory neuron clusters were obtained and merged into a single cluster for the remaining analysis. In addition, clusters for oligodendrocyte precursor cells and oligodendrocytes were merged due to the low number of cells.

### scRNA-seq data analysis

Adult mouse primary visual cortex scRNA-seq data(*47*) were obtained from NCBI GEO (accession GSE71585) with the annotation and processed TPM files. We used the cell-type annotations from the original study, representing 10 major cell types. For the excitatory neuron expression profile, we used layer 2/3 excitatory neurons. We compared the z-score normalized gene expression profiles of 56 genes that were commonly profiled by scRNA-seq and RNA seqFISH. The degree of similarity was evaluated by using the Pearson correlation.

### 3D visualization of the data

DNA seqFISH+ data and immunofluorescence signals were visualized in 3D using PyMOL (Molecular Graphics System, v.2.0 Schrödinger) as described previously(*32*). DNA seqFISH+ data were visualized by generating a .xyz file containing the x, y and z coordinates of each FISH probe coordinate. Each FISH probe coordinate was displayed as a sphere, and sticks were drawn between genomically adjacent coordinates for some of the visualization. Immunofluorescence signals were visualized by displaying a surface around x, y and z coordinates with intensity z-score values above 2 from raw images or with the binarized images from the foci detection.

### Estimation for DNA seqFISH+ detection efficiency

The detection efficiency of DNA seqFISH+ for the post-mitotic diploid cells in the female mouse brain cortex was estimated as follows. The total number of expected FISH spots for 3,660 loci in the autosomal or X chromosomes is 7,320 in the female diploid cells. In our DNA seqFISH+ experiments, we observed 2,813.0 ± 1,334.0 (median ± s.d.) and 3,884.0 ± 1,252.9 (median ± s.d.) spots per cell for all 2,762 cells and a subset of 701 cells found at the center z sections, respectively, and their detection efficiencies can be estimated as 38.4 ± 18.2% (median ± s.d.) and 53.1 ± 17.1% (median ± s.d.), showing the similar detection efficiency with DNA seqFISH+ experiments in mouse ES cells(*32*).

### Spatial separation of homologous chromosomes

Both for the 1-Mb and 25-kb resolution DNA seqFISH+ data, the whole homologous chromosomes or chromosomal regions were separated by the DBSCAN clustering algorithm in scikit-learn library in Python as performed previously(*32*). In the further downstream analysis that required homologous chromosome separation, only chromosomes with two homologous chromosomes detected in a cell were used for the 1-Mb data while chromosomes with at least one chromosomal region detected in a cell were used for the 25-kb data. The lack of two homologous chromosome detection could be due to the spatial intermingling of homologous chromosomes, which typically happened with the 1-Mb resolution data, or incomplete coverage of the nucleus in the z sections of the images.

To separate the Xa and Xi, z-score normalized mean Xist intensities per separated homologous chromosomes across detected loci were first calculated. Then the z-score normalized mean Xist intensities were thresholded with z-score equal or below 1 for Xa and z-score above 1 for Xi. This gave 980 Xa and 936 Xi (51.1% vs. 48.9%) in the 1-Mb resolution data and 2,019 Xa and 1,956 Xi (50.8% vs. 49.2%) in the 25-kb resolution data from 2,762 cells in the three biological replicates.

### Spatial distance versus genomic distance analysis

The 3D coordinates with μm units (x: 0.103, y: 0.103, z: 0.250 μm per voxel) were used to compute the spatial distance of pairs of loci from DNA seqFISH+ data. To calculate the median spatial distance from given pairs of loci within a given homologous chromosome or chromosomal region across all cells or cells in each cell type for the 1-Mb and 25-kb resolution data, we calculated the Euclidean distances between all pairs of detected loci within only homologous chromosomes that were spatially separated in each cell, and tabulated them with their genomic distances. Similarly, to calculate the quartile spatial distance from given pairs of loci within chromosomes across all cells or cells in each cell type for the 1-Mb resolution data, we calculated the Euclidean distances between all pairs of detected loci within individual chromosomes without separating homologous chromosomes in each cell, and tabulated them with their genomic distances. For the 1-Mb resolution data, both median spatial distances within homologous chromosomes and quartile spatial distances within chromosomes in each cell were highly correlated. Those spatial distance maps in each chromosome were compared with the Hi-C maps using Spearman correlation. The spatial distance of inter-chromosomal loci was similarly computed by calculating the quartile spatial distance from given pairs of inter-chromosomal loci across cells in each cell type for the 1-Mb resolution data.

To compute the relationships between spatial distances between pairs of loci versus genomic distances, the pairs of genomic loci were grouped at given genomic bins, and median distances from each genomic bin were computed using quartile spatial distance within chromosomes for the 1-Mb resolution data and median spatial distance within homologous chromosomal regions for the 25-kb resolution data.

### DNA spatial proximity map analysis

To generate pairwise spatial proximity maps from the DNA seqFISH+ dataset, the spatial distance of pairs of loci were first computed as described above. Then the fractions of pairs of loci within a certain radius were computed. For the 1-Mb resolution data, we used all detected pairs of loci within individual cells, while for the 25-kb resolution data, we used detected pairs of loci within individual chromosomal regions in each homologous chromosome. We used a search radius of 500 nm for the 1-Mb resolution data and 150 nm for the 25-kb data based on the previous studies(*22, 27, 30, 32*). The pairwise spatial proximity maps were compared with the Hi-C maps using Pearson correlation.

### Radial positioning analysis

The segmented nuclei were first individually normalized using Euclidean distance transformation from their centroids using bwdist function in MATLAB, and then radial distances were scaled from 0 (nuclear center) to 1 (nuclear periphery), similarly to those previously described(*101, 102*). To investigate the radial positioning of the genomic loci, the radial scores at the rounded DNA seqFISH+ spot voxels were obtained from individual nuclei. The median radial positioning of loci in each cell type was then compared with the chromosome size, chromosome-wide gene density(*73*) and SF3a66 chromatin profiles.

### DAPI meta feature analysis

Meta feature extraction from DAPI images was performed based on the previous studies for cell optical phenotype extraction in the fluorescence images(*103, 104*) with modifications. In brief, the following 16 imaging features were extracted from the DAPI images for each segmented nucleus using regionprops3 function in MATLAB, including volume, convex volume, surface area, three principal axis lengths, extent, solidity, total intensity, mean intensity, median intensity, standard deviation of intensity, lower quartile intensity, upper quartile intensity, skewness of intensity, kurtosis of intensity. Then each feature was normalized by computing z-score per each biological replicate. After combining cells from three biological replicates, cells were clustered based on the DAPI meta features with hierarchical clustering, visualized with UMAP(*100*) using a UMAP-learn library in Python, and compared to transcriptionally defined cell types. For this analysis, we used a subset of 701 cells from three biological replicates at the center z sections in each image to eliminate the volume differences due to the incomplete coverage of nuclei in z sections.

### Global chromatin state analysis

The averaged intensity for individual immunofluorescence markers in each segmented nucleus was first normalized by mean intensity of the marker from nuclei in each field of view to correct the intensity bias among fields of view and different replicates. After combining all the nuclei (n = 2,762 cells) from three biological replicates, the normalized intensity for each marker was further normalized by computing z-score. The normalized intensity profiles from 8 immunofluorescence markers for each cell were used to compare cell-type-specific global chromatin states as histograms and visualize individual cells in a reduced dimension with UMAP(*100*) using a UMAP-learn library in Python.

### Nuclear foci analysis

The processed foci from DAPI, MajSat, SF3a66, ITS1 and Xist images, corresponding to fluorescence intensity enriched voxels for each marker described under *‘Nuclear marker image analysis’*, were quantified to measure foci number per cell, individual foci volume (μm), total foci volume per cell (μm^3^), and radial position distribution at their centroids for each cell type. To obtain the radial positioning of H3K27me3 globules that do not correspond to the Xist foci, voxels with raw intensity z-score above 2 and Xist raw intensity z-score 2 or below were used for each cell type. Similar to the DAPI meta feature analysis above, we used a subset of 701 cells from three biological replicates at the center z sections in each image to eliminate the foci property differences originating from the incomplete coverage of nuclei in z sections.

### Chromatin profile and fixed point analysis

The imaging-based chromatin profile analysis was performed using sequential immunofluorescence and 1-Mb resolution DNA seqFISH+ data as described previously(*32*). In brief, we calculated the spatial distances between each DNA locus and the nearest exterior of “hot” immunofluorescence voxel, defined by two standard deviations above the mean value for the edge processed each immunofluorescence intensity (described under ‘*Nuclear marker image analysis’*) in each nucleus. From this distance metric, we generated a “chromatin profile” by counting the percentage of cells in which each DNA locus is within 300 nm of the exterior of each immunofluorescence mark, the resolution of the diffraction-limited immunofluorescence images. The chromatin profiles among brain cell types and mouse ES cells(*32*) were compared with Pearson correlation. The chromatin profiles in each cell type were further compared to 1-Mb resolution gene density and radial positioning of loci with Pearson correlation and Spearman correlation. Using the chromatin profiles, fixed points were determined as loci that appear 2 standard deviations above the mean percentage score for each immunofluorescence mark in autosomes.

### Single chromosome domain analysis

#### Normalization of pairwise spatial distance

For each assigned allele ID (Xist state ID for X chromosome), we independently computed the Euclidean distance (in μm) between pairs of DNA loci, which were mapped to the reference locus IDs. Euclidean distances were set as NA for pairs of loci with at least one being undetected. Under rare occasions, in the same single cell allele when more than one imaged loci were mapped to the same locus ID, we took the median of all available pairwise Euclidean distances. Chromatin contacts within each single cell allele were represented as a separate distance matrix.

The raw Euclidean distances were normalized by expected values from genomic distances, to adjust for the basal level of interactions between adjacent loci in the linear genome(*30*). Briefly, between every pair of regions, we computed the genomic distance in kilobases for 25-kb resolution data and the median spatial Euclidean distances over all detected single cell alleles. We then constructed a local polynomial regression (loess) model between genomic and ensemble spatial distances, using the stats::loess() function with default parameters in R. For every pair of genomic loci i and j, the estimated spatial distance was predicted from the local polynomial model, and the normalized distances were calculated by:

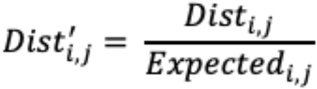

Here *Dist_i,j_* represents the raw spatial distance between loci i, j, and Expected_i, j_ is the expected distance predicted using the loess model as mentioned above.

#### Proximity measurements between genomic loci

Given that we were interested in loci with close proximities, we implemented a Gaussian kernel to convert spatial distances into a fixed range between 0 and 1, where loci with high spatial distances received proximity scores of zero and would not contribute to the variance characterization steps. The transformation into proximity score K_i,j_ is specified by the formula below, with Dist’_i,j_ being the normalized distance from linear genome:

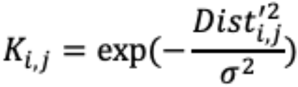

The band width parameter *σ* was chosen such that the proximity between genomic loci decreased exponentially and approached zero with large spatial distances. From the imaging results, we found that 0.4 *μ*m served as a cutoff distance for loci interaction, which we would like to use as the criteria for choosing ***ό***. By performing a grid search and comparing the resulting proximity scores versus raw spatial distance, we found that at ***ό*** = 0.4, 0.4 *μ*m corresponded to an average proximity score of 0.5, and at 1.5 *μ*m for all loci the proximity scores reached zero.

We also noticed that the presence of missing data could interfere with the quantitative characterization of heterogeneity between single cell alleles. Therefore, we implemented a simple smoothing approach to the proximity matrices, where missing values were substituted by the mean of neighboring entries (pooling from entries within 1 unit of row/column indices). If missing data still existed after smoothing, they were assigned zeroes in the proximity matrix.

#### Characterization of major variations

Since single chromosomes in the 25-kb resolution data were represented as symmetric two-dimensional proximity matrices, we vectorized each matrix by ‘rolling-out’ the upper triangle, and concatenated the vectorized proximity scores. In the new concatenated data, each row was the pairwise proximity score and each column represented one allele in a cell. Furthermore, for each chromosome we selected only cells with two identifiable copies available, and with both alleles below a predefined threshold of missing data frequencies (80%) in raw distance matrices. To adjust for bias resulting from overall packing level and missing data frequencies, the concatenated matrices were standardized and scaled using the Seurat::ScaleData() function(*105*), with average raw pairwise distances and total number of detected regions as latent variables to regress out.

We then used principal component analysis (PCA) to characterize the variations of proximities in single cell alleles, using a set of most variable features (pairwise proximities) based on standardized variances implemented in Seurat::FindVariableFeatures()(*106*). We selected interactions with the top 70% highest variance as the most variable features. Additionally, we also filtered input features by the frequency of being undetected across single cell alleles, with a threshold of < 70% for 25kb data. PCA was performed using the Seurat::RunPCA() function on the concatenated variable features by alleles matrix for individual chromosomes.

#### Heterogeneity of pairwise spatial proximities

We projected PCA results onto all features by computing the dot product between scaled proximities and allele embeddings (rotated allele coordinates in PC space). The signs and absolute values of projected feature loadings indicated the dimensions of variations identified by each PC. By examining projected feature loadings and the approximate singular values of PCs for each chromosome in elbow plots, we chose to use the top 10 or 20 PCs to construct a shared nearest neighbor graph for Leiden clustering, using the function Seurat::FindNeighbors() and Seurat::FindClusters()(*107, 108*). The resolution setting was 0.8, so as to return 10-20 structural clusters. To examine cluster distribution and pairwise spatial proximity patterns, we used UMAP(*100*) for visualization, and further computed the median of pairwise spatial distance for every cluster to summarize cluster profiles.

#### Boundary score analysis

We also calculated boundary scores to characterize possible domain boundaries of different clusters. Similar to the previous study(*23*), we took the 4 (or 8 for ChrX) loci upstream and downstream of every target genomic locus (including itself) as two neighboring ‘domains’, forming two domains of sizes 5 (or 9 for ChrX) including the target locus. We then calculated and calculated the inter- and intra-domain distances between all detected loci, using genomically normalized distances. The raw boundary score was defined as the division of the median of inter-domain distances by the median of intra-domain distances. Accordingly, we computed boundary scores for single cell alleles, and further normalize the boundary scores for all loci in single allele (x) to values between 0 and 1:

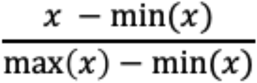

We then computed the median of each genomic locus for alleles from the same cluster, in order to derive a cluster-level boundary score matrix for every chromosome.

## Supplementary Figures

**Fig. S1.**
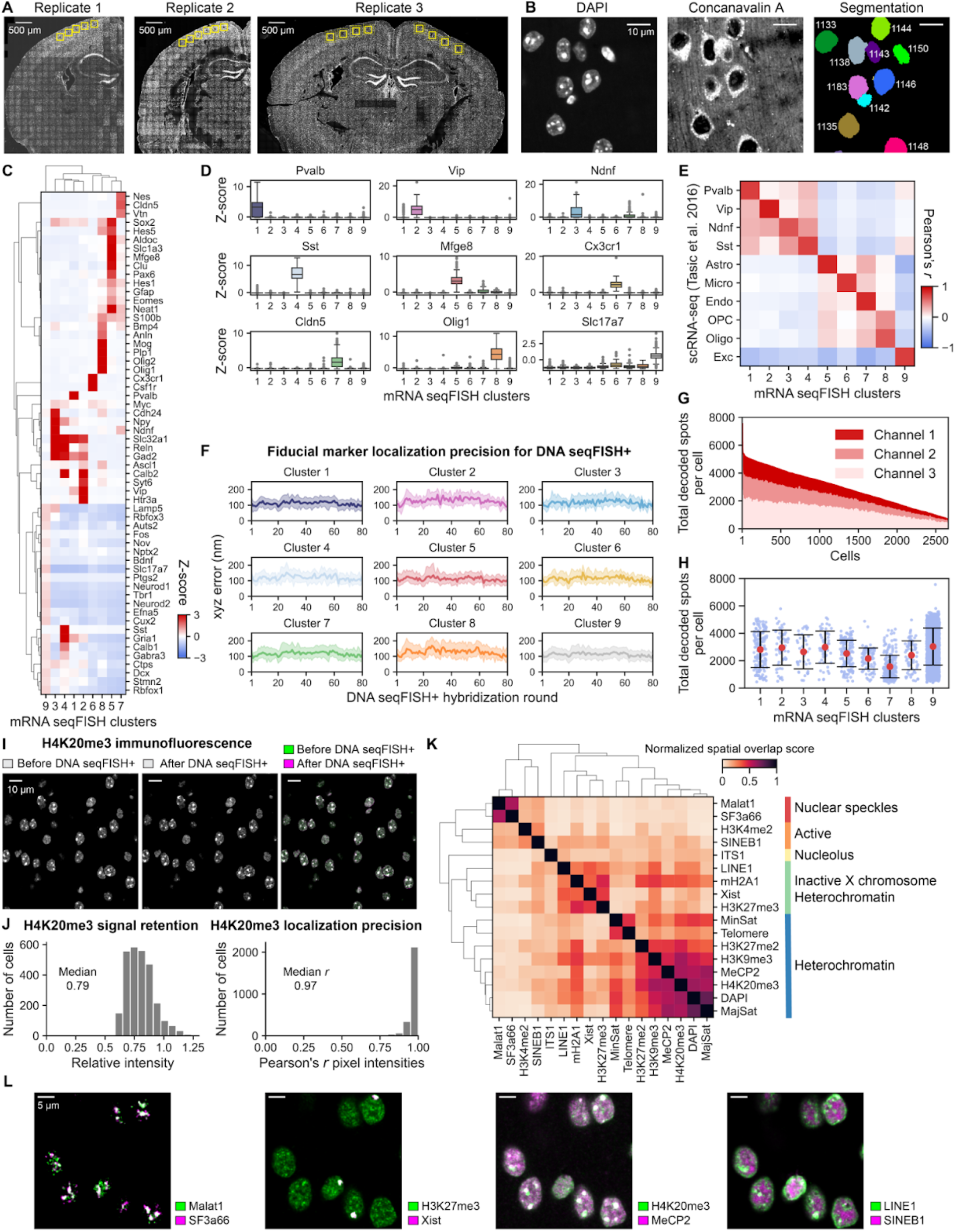
Validation for the integrated spatial genomics approach in mouse brain cortex. (A) DAPI images of the mouse brain section with three biological replicates. Yellow boxes represent the imaged fields of view in each replicate. (B) Representative images for DAPI and concanavalin A staining for nuclear segmentation, and segmented nuclei in the mouse cortex. (C) The hierarchical clustering of z-scored mRNA expression profiles by RNA seqFISH. The gene expression profiles for each RNA seqFISH cluster are distinct. (D) Box plots showing z-scored mRNA expression levels for cells in each RNA seqFISH cluster. The box plots represent the median, interquartile ranges, whiskers within 1.5 times the interquartile range, and outliers. (E) The transcriptionally defined cell clusters by RNA seqFISH correlate well with previously reported cell types by scRNA-seq in mouse cortex(*47*). Based on the matches between RNA seqFISH cluster and scRNA-seq cell type represented by the highest Pearson’s correlation coefficient, RNA seqFISH clusters were annotated. (F) Quantification of the fiducial marker localization precision in RNA seqFISH cluster for 80 hybridization rounds in the DNA seqFISH+ experiments, relative to the reference fiducial marker image. Shaded regions represent standard deviation. (G) The total number of DNA seqFISH+ spots detected in each of the fluorescent channels in single cells from all cell types. Fluorescent channels 1 and 2 contain the 1-Mb resolution data and channel 3 contains the 25-kb resolution data. (H) The total number of DNA seqFISH+ spots detected in each cell type. (I) Preservation of the nuclear structure with the integrated spatial genomics protocol for tissue sections. Good colocalization (white in the right panel) of H4K20me3 immunofluorescence signals before and after DNA seqFISH+ preparation including a heating denaturation step. (J) Quantification of the H4K20me3 immunofluorescence signal retention in the nuclei before and after DNA seqFISH+ preparation (left) and localization precision measured by Pearson correlation of pixel intensities with a single z-section (right). (K) Quantification of spatial overlap of voxels with intensity z-score above 2 between pairs of nuclear marker images stained by sequential immunofluorescence, RNA FISH, or DNA FISH. (L) Representative images of nuclear markers in (K), showing either colocalization with white color (Malat1 and SF3a66; H3K27me3 and Xist; H4K20me3 and Mecp2) or mutually exclusive localization pattern (LINE1 and SINEB1). n = 2,762 cells from three biological replicates were used for quantification in (C)-(K).

**Fig. S2.**
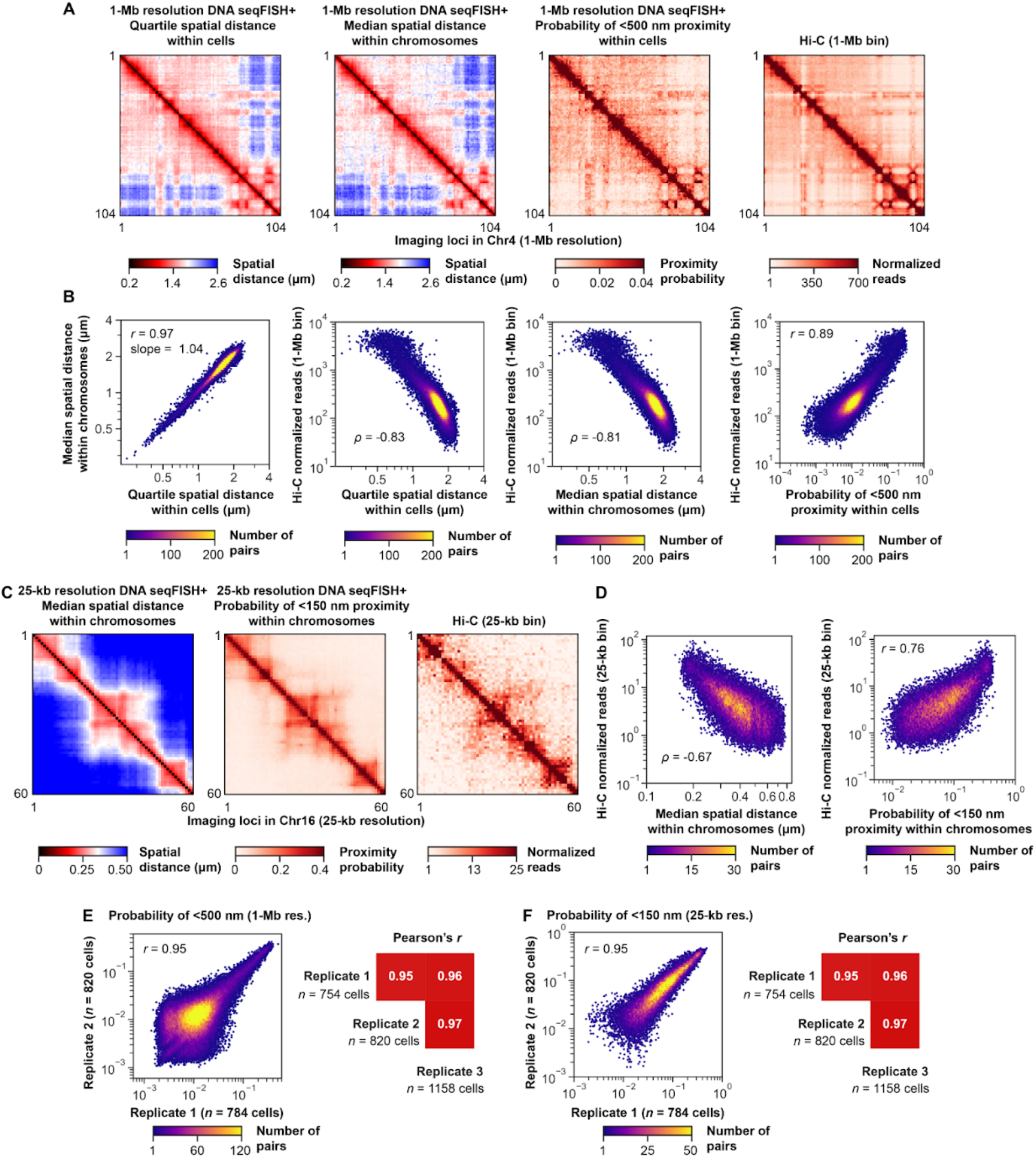
Validation for DNA seqFISH+ in mouse brain cortex. (A) Heat maps of 1-Mb resolution chromosome 4 showing quartile spatial distances of pairs of loci within cells, median spatial distances of pairs of loci within homologous chromosomes, probabilities of pairs of loci within a search radius of 500 nm within cells by DNA seqFISH+, and the corresponding Hi-C map(*6, 56*) with 1-Mb resolution binning, respectively (left to right). (B) Comparison between spatial distances (left), between spatial distances and Hi-C dataset (middle panels), and between spatial proximity probabilities and Hi-C dataset (right) with pairs of intra-chromosomal loci in all autosomes with 1-Mb resolution. (C) Heat maps of 25-kb resolution chromosome 16, showing median spatial distances of pairs of loci within homologous chromosomes (left), probabilities of pairs of loci within a search radius of 150 nm within cells by DNA seqFISH+, and the corresponding Hi-C map(*6, 56*) with 25-kb resolution binning. (D) Comparison between spatial distances and Hi-C dataset (left), and between spatial proximity probabilities and Hi-C dataset (right) with pairs of intra-chromosomal loci in all autosomes with 25-kb resolution. n = 2,762 cells from three biological replicates were used for DNA seqFISH+ quantification in (A)-(D). (E and F) Pearson’s correlation of probabilities for the pairs of loci within a search radius of 500 nm (1-Mb resolution data) and 150 nm (25-kb resolution data) among three biological replicates of DNA seqFISH+ experiments. All unique intra-chromosomal pairs of loci were calculated for the 1-Mb (n = 2,460 loci) and 25-kb (n = 1,200 loci) resolution data with n = 754, 820, and 1,158 cells for biological replicates 1, 2, and 3, respectively. r and ρ represent Pearson’s correlation coefficient and Spearman’s correlation coefficient in (B), (D), (E), (F).

**Fig. S3.**
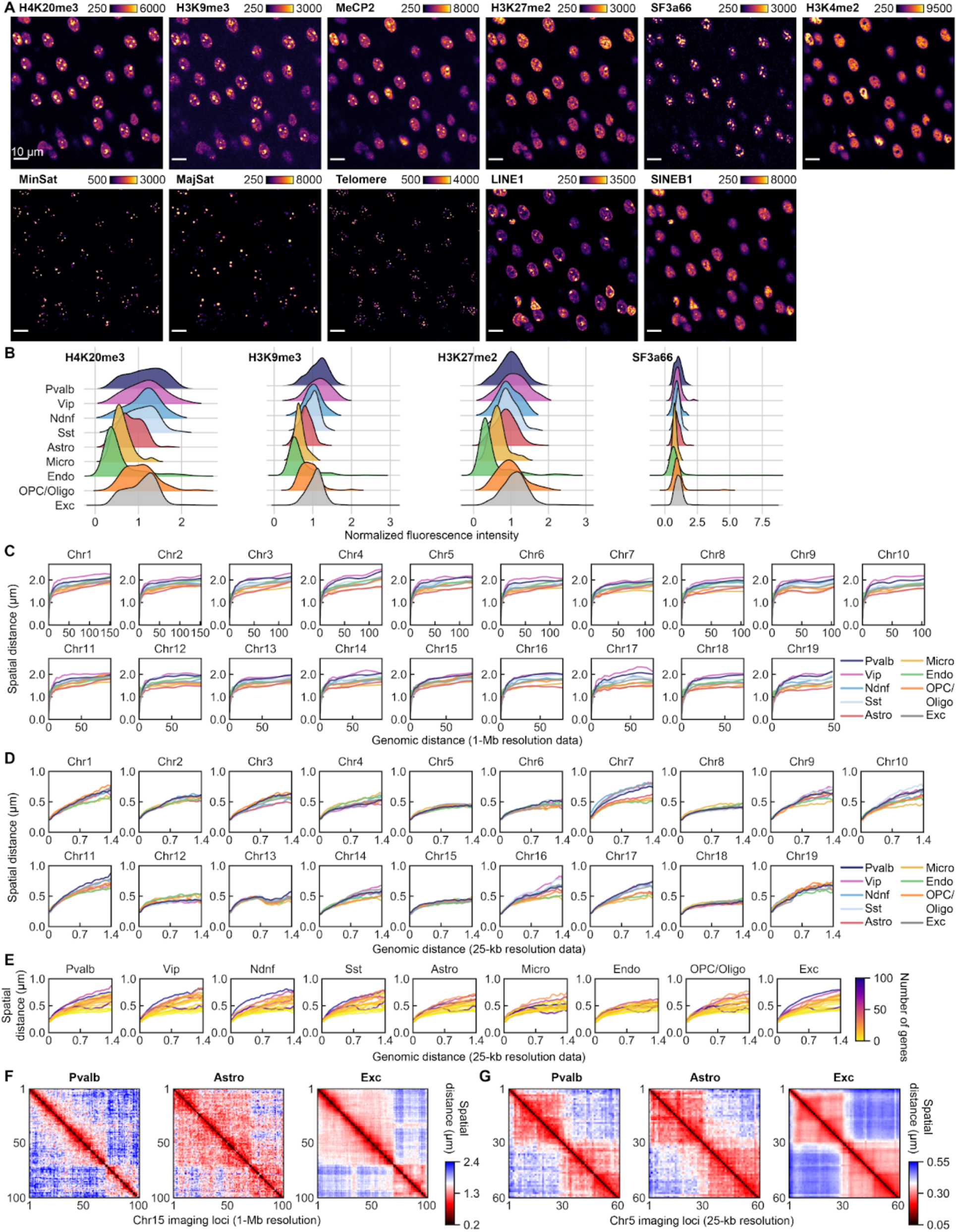
Cell-type-specific global chromatin states and chromosome scaling analysis. (A) Sequential immunofluorescence staining (top) and DNA FISH (bottom) images from a single z-section of the mouse cortex. Corresponding cell types of the images are shown in Fig. 2A. Color bars represent fluorescence intensity (a.u.). Scale bars, 10 μm. (B) Kernel density estimations of normalized IF intensities show cell type dependent distributions of marker levels. (C and D) Physical distance as a function of genomic distance across transcriptionally defined cell types at the 1-Mb resolution in (C) and at 25-kb resolution in (D) for each autosome. Similar plots for the active and inactive X chromosome are shown in fig. S8G. (E) Physical distance as a function of genomic distance comparing different 25-kb resolution chromosomal regions in each cell type, colored by the total number of genes in each chromosomal region. n = 2,762 cells from three biological replicates for quantification in (B)-(E). (F and G) Spatial distance between pairs of intra-chromosomal loci within cells by DNA seqFISH+ in different cell types at 1-Mb resolution (quartile spatial distance within cells for chromosome 15) in (F) and at 25-kb resolution (median spatial distance within homologous chromosomes for chromosome 5) in (G). n = 1,895, 155, and 152 cells for excitatory neurons, Pvalb inhibitory neurons, and astrocytes from three biological replicates in (F), (G).

**Fig. S4.**
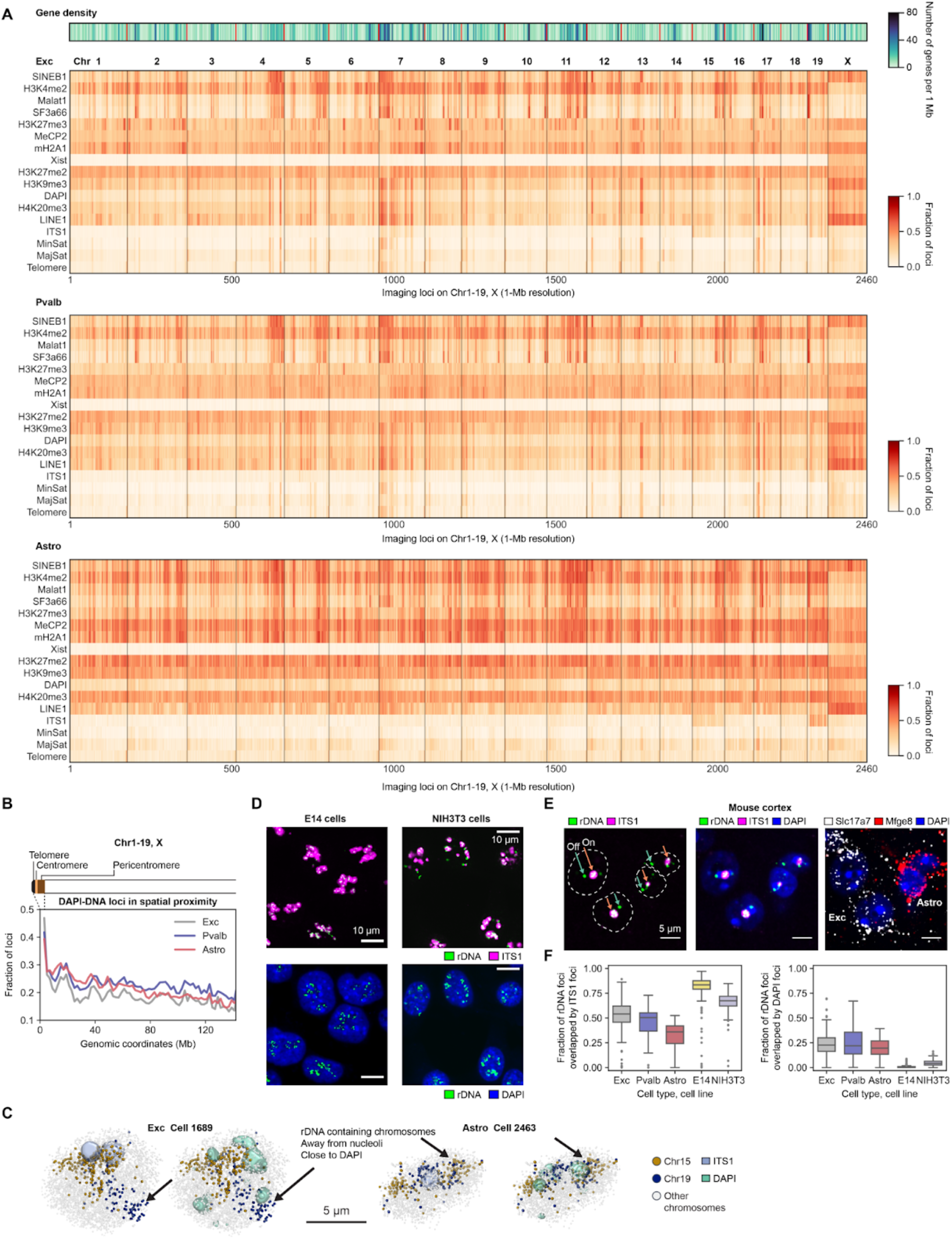
Cell-type-specific chromatin profiles and rDNA association with nucleoli. (A) Cell type specific chromatin profiles for 1-Mb resolution DNA seqFISH+ loci (n = 2,460 loci), calculated by the fraction of loci found within 300 nm of each subnuclear marker exterior. Gene density in each 1-Mb resolution loci is shown at the top. (B) Cell type specific fraction of loci from DAPI exterior for 1-Mb resolution DNA seqFISH+ loci from 20 chromosomes. The genomic loci were grouped based on their genomic coordinates in each chromosome. The loci that are genomically close to the pericentromeric repetitive region, a component of constitutive heterochromatin, showed higher spatial association with DAPI-enriched constitutive heterochromatic bodies, consistent with the observation in mouse ES cells(*10, 32*). n = 1,895, 155, and 152 cells for excitatory neurons, Pvalb inhibitory neurons, and astrocytes from three biological replicates for quantification in (A) and (B). (C) Representative 3D images of nucleoli by ITS1 RNA FISH, heterochromatic bodies stained by DAPI, and chromosomes 15 and 19 by DNA seqFISH+ in the excitatory neuron (left) and astrocyte (right). Despite containing rDNA regions(*69*), those chromosomes did not necessarily associate with the nucleolus in some of the cells (Cell 1689, 2463 are shown as examples). (D) rDNA staining by DNA FISH with nucleolar imaging by ITS1 RNA FISH and DAPI staining in E14 mouse ES cells (left) and NIH3T3 fibroblast cells (right) from a single z-section. Scale bars represent 10 μm. (E) Unlike the cultured cells in (D), rDNA staining confirms that some of the ribosomal repeat regions appear outside of the nucleolus and locate at DAPI-enriched heterochromatin in post-mitotic cells in the brain. The images are from a single z-section and scale bars represent 5 μm. (F) The fraction of rDNA loci present in the nucleolus or close to heterochromatic regions is cell type dependent with significant off-nucleolus association in the brain compared to cultured mouse ES cells or fibroblasts. The boxplots represent the median, interquartile ranges, whiskers within 1.5 times the interquartile range, and outliers. n = 365, 18, 32 for excitatory neurons, Pvalb inhibitory neurons, and astrocytes from 5 fields of view in one mouse brain section, and n = 305, 334 cells for E14 mouse ES cells and NIH3T3 cells from 7 and 8 fields of view in each sample.

**Fig. S5.**
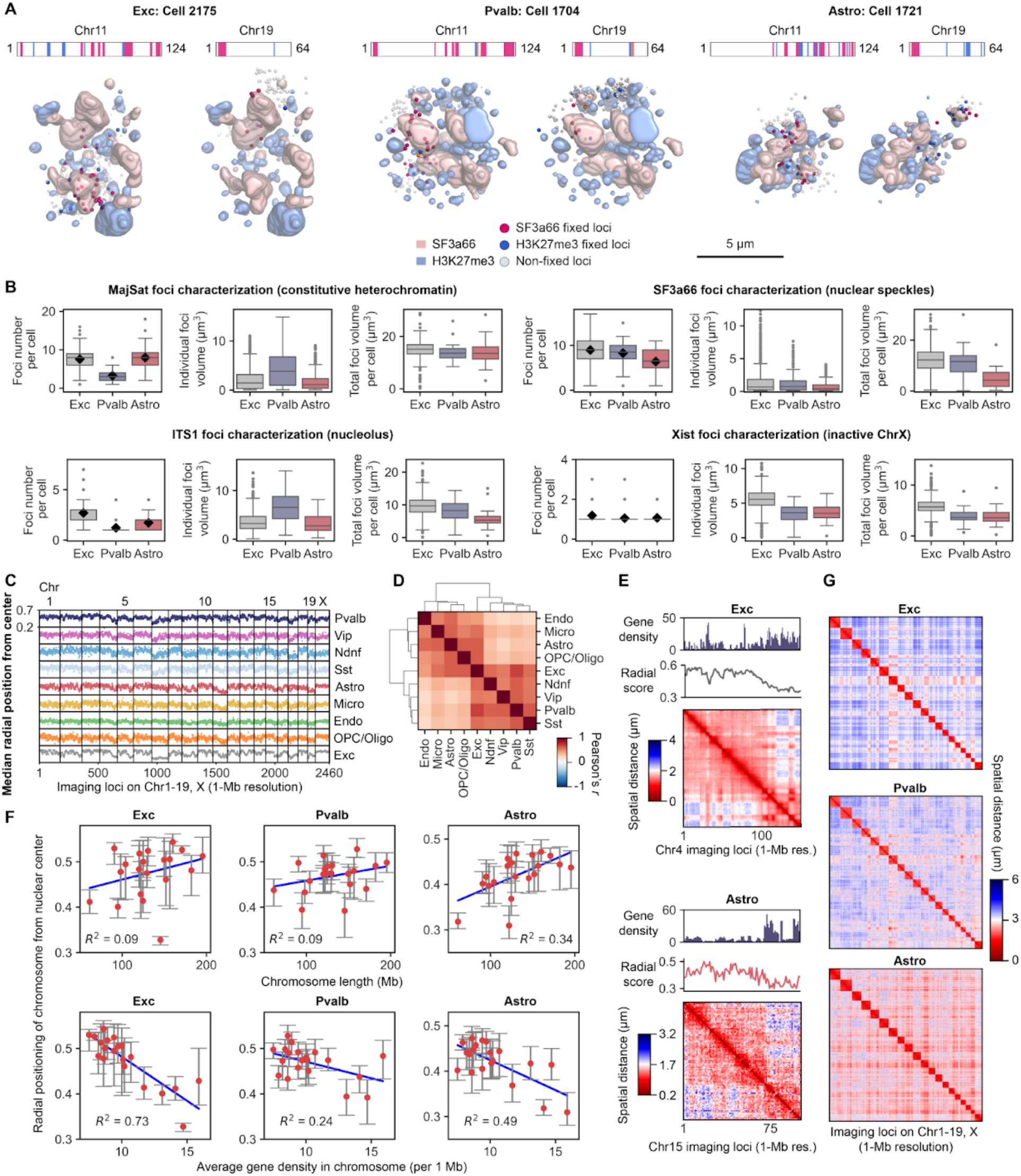
Cell-type-specific nuclear foci and radial chromosome organization, and common nuclear organization. (A) Images of individual chromosomes (Chr11 or Chr19) and markers (SF3a66 and H3K27me3), showing each chromosome has cell-type specific fixed points and span the corresponding nuclear bodies in single cells. (B) Characterization of subnuclear marker foci features across 3 major cell types, representing cell-type specific nuclear foci number and volume distributions. The boxplots represent the median, interquartile ranges, whiskers within 1.5 times the interquartile range, and outliers, and the black diamond dots represent mean. Some outliers were omitted at the display range for visual clarity (6 outliers in the middle panel of the SF3a66 foci characterization). n = 460, 33, and 48 cells for excitatory neurons, Pvalb inhibitory neurons, and astrocytes in the center z-sections from three biological replicates. (C) Median radial positioning from the nuclear center for all 2,460 loci imaged at 1-Mb resolution for all cell types. (D) Hierarchical clustering of radial positioning profiles from (C) between pairs of cell types, showing neuronal and glial cell clusters. n = 2,762 cells from three biological replicates for quantification in (C) and (D). (E) Examples of the comparison among gene density, radial positioning of loci, and spatial distance of pairs of loci in excitatory neurons (top) and astrocytes (bottom). Gene dense large-scale domains with tens of megabases in size tend to localize in the nuclear interior in both cell types. Similar observation about consistency between large-scale domain radial positioning and large-scale chromosome conformation capture patterns was made by the single-cell chromosome conformation capture study(*18*). (F) Radial nuclear positioning for 20 chromosomes showed cell-type-specific dependency on chromosome length (top) and chromosome-wide gene density (bottom). The red dots represent median for individual chromosomes and error bars represent interquartile ranges. The coefficient of determination for linear regression is shown in each plot. (G) Cell-type-specific quartile spatial distance between pairs of inter-chromosomal loci (n = 2,460 loci for 20 chromosomes) in three major cell types. n = 1,895, 155, and 152 cells for excitatory neurons, Pvalb inhibitory neurons, and astrocytes from three biological replicates for quantification in (E)-(G).

**Fig. S6.**
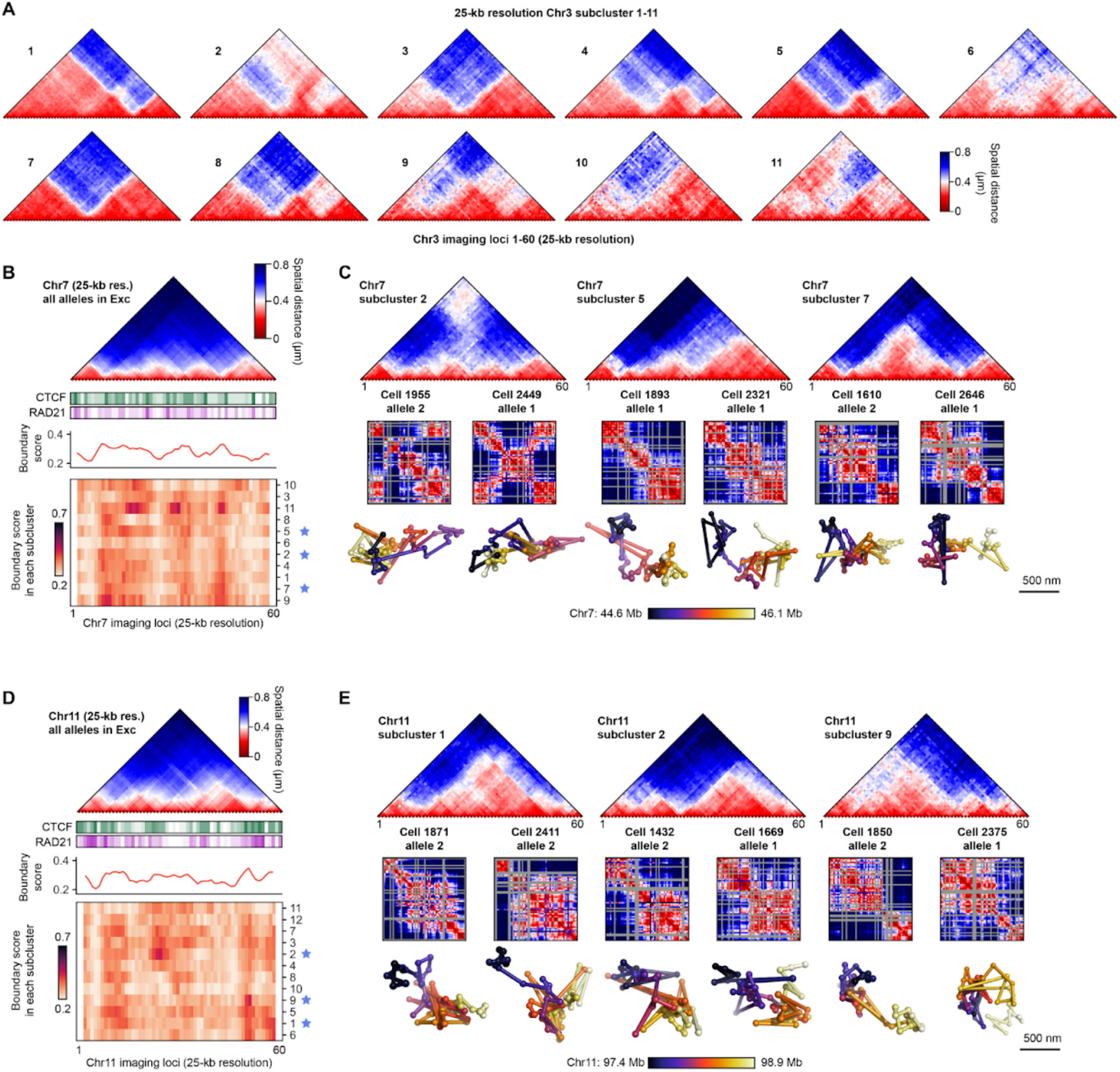
Heterogeneous domain structures in single chromosomes without clear ensemble domain structures. (A) Pairwise spatial distance maps from all 11 structural subclusters of the 25-kb resolution data in chromosome 3 (n = 1,524 chromosomes) from excitatory neurons. Bulk averaged domain structure for the chromosome 3 region is shown in Fig. 6D. (B) Bulk averaged pairwise spatial distance map from excitatory neurons (top) with CTCF and cohesin binding sites(*77*). Single chromosomes were clustered into different domain structures with distinct median boundary score profiles (bottom) for the chromosome 7 region (44.6-46.1 Mb; n = 1,386 chromosomes). Despite the lack of clear domains at the bulk level, there are several domain subclusters with different configurations in single chromosomes. (C) Examples for three of the structural subclusters are shown with different pairwise spatial distance map structures (top). Corresponding single chromosome distance maps (middle) and 3D chromosome structures (bottom) are shown. (D and E) Similar to (B and C) for the chromosome 11 region (97.4-96.9 Mb) (n = 1,514 chromosomes). Chromosomes with >20% loci detected in n = 1,895 excitatory neurons from three biological replicates were used for analysis in (A)-(E).

**Fig. S7.**
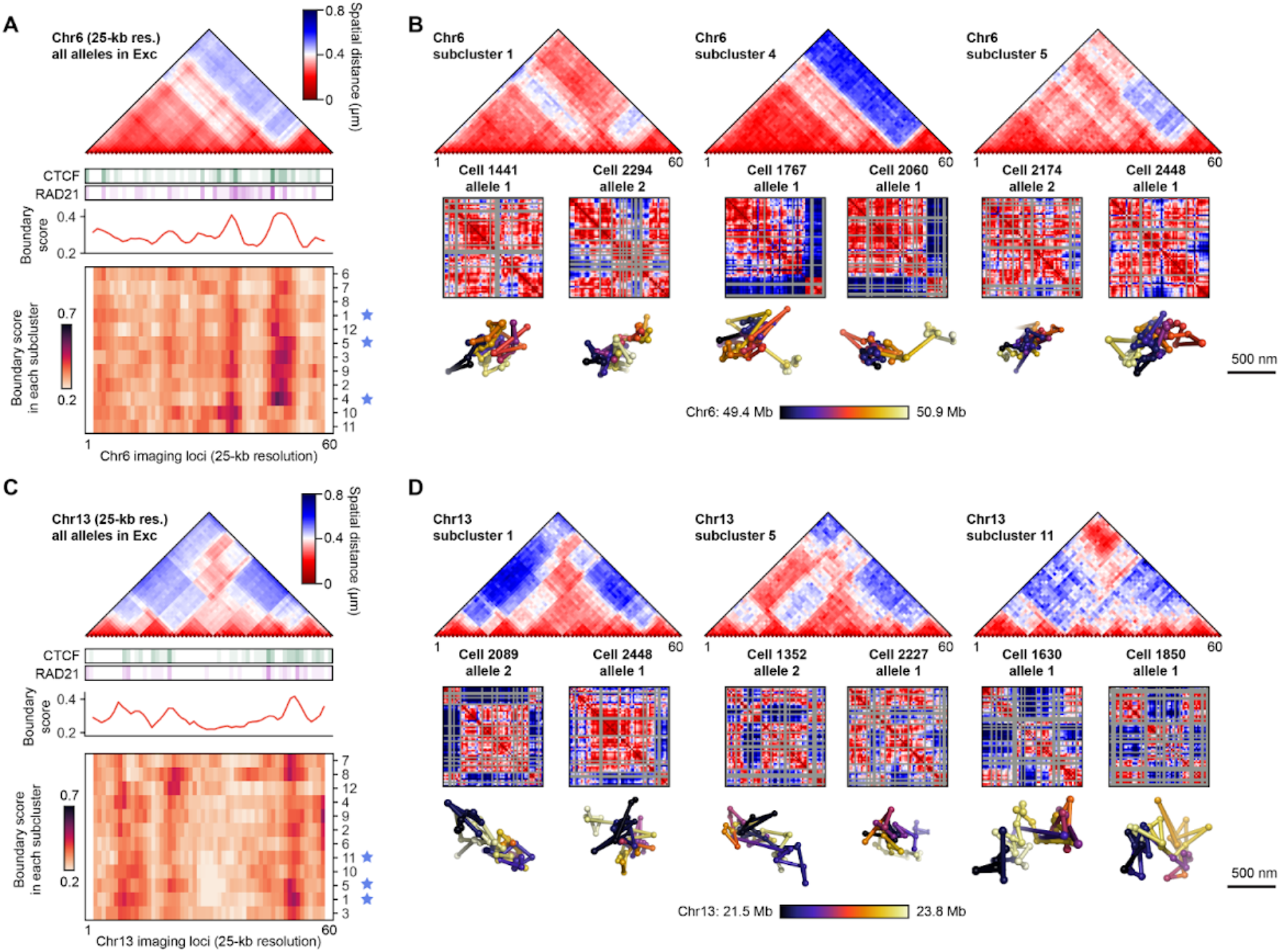
Heterogeneous higher-order domain structures in single chromosomes. (A) Bulk averaged pairwise spatial distance map from excitatory neurons (top) with CTCF and cohesin binding sites(*77*). Single chromosomes were clustered into different domain structures with distinct median boundary score profiles (bottom) for the chromosome 6 region (49.4-50.9 Mb; n = 1,550 chromosomes). (B) Examples for three of the structural subclusters are shown with different pairwise spatial distance map structures and higher-order domain structures (top). Corresponding single chromosome distance maps (middle) and 3D chromosome structures (bottom) are shown. (C and D) Similar to (A and B) for the chromosome 13 region (21.5-23.8 Mb; n = 1,414 chromosomes). Chromosomes with >20% loci detected in n = 1,895 excitatory neurons from three biological replicates were used for analysis in (A)-(D).

**Fig. S8.**
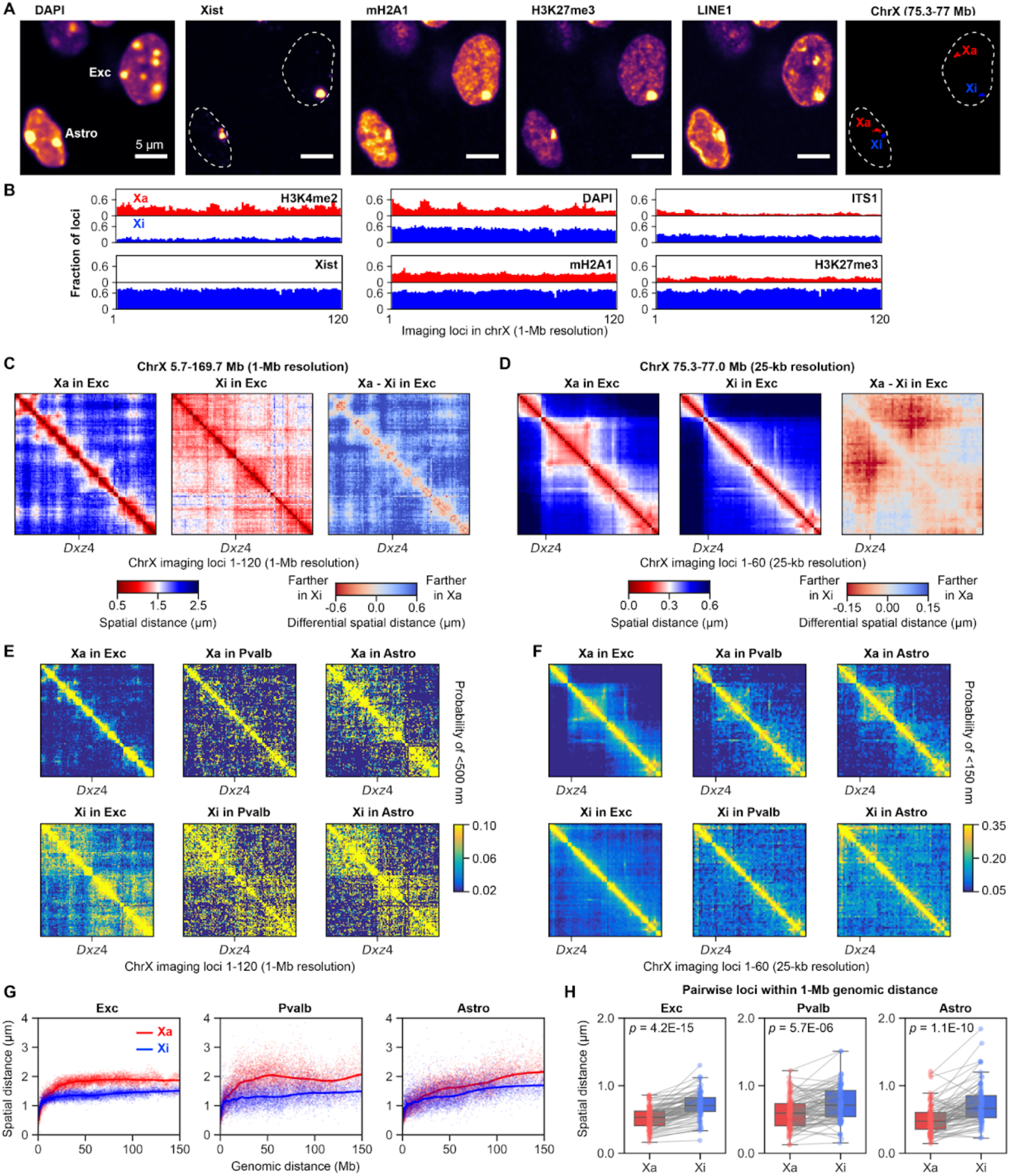
Active and inactive X chromosome organization across cell types. (A) DAPI, Xist RNA and IF staining distinguishes the active X chromosome (Xa) and inactive X chromosome (Xi) in single cells in the mouse brain cortex. The images for DAPI, Xist RNA and IF staining are the maximum intensity z-projected from 1.5 μm z-slices, and the image for Xa and Xi is decoded 25-kb resolution DNA seqFISH+ data from all z-slices. The scale bars represent 5 μm. (B) Chromatin profiles of the Xa and Xi with respect to the H3K4me2, Xist, DAPI, mH2A1, ITS1 and H3K27me3 across 120 loci at 1-Mb resolution in excitatory neurons. (C) Pairwise median distance maps for Xa (left) and Xi (middle) in excitatory neurons from the 1-Mb DNA seqFISH+ data, and differential distance map between Xa and Xi (right). Xi is more compact at long genomic distances, but locally is less structured than Xa at the finer scale. The spatial distance maps were computed from n = 1,895 excitatory neurons with spatially separated Xa or Xi detected from three biological replicates. (D) Pairwise median distance maps for Xa (left) and Xi (middle) in excitatory neurons from the 25-kb DNA seqFISH+ data. Differential distance map (right) shows more compact averaged structure at the finer resolution for Xa. The spatial distance maps were computed from n = 1,895 excitatory neurons with at least one Xa or Xi region (75.3-77.0 Mb) detected in a cell from three biological replicates. (E) Heat maps for probabilities of pairs of loci within a search radius of 500 nm in Xa (top) and Xi (bottom) by 1-Mb resolution DNA seqFISH+ in three major cell types. (F) Heat maps for probabilities of pairs of loci within a search radius of 150 nm in Xa (top) and Xi (bottom) by 25-kb resolution DNA seqFISH+ in three major cell types. (G) Physical distance as a function of genomic distance for Xa (red) and Xi (blue) in three major cell types by 1-Mb resolution DNA seqFISH+. Color dots represent each pair of genomic loci, and colored lines represent median spatial distance of pairs of genomic loci within genomic distance bins. (H) Box plots with spatial distance for pairs of genomic loci within 1-Mb genomic distance in 1-Mb resolution data in the Xa and Xi in three major cell types, overlaid with individual pairs of genomic loci. Contrary to the larger-scale genomic distance in (G), at the finer scale below 1 Mb, Xa had smaller spatial distances between pairs of genomic loci compared to Xi. The boxplots represent the median, interquartile ranges, whiskers within 1.5 times the interquartile range, and p values were calculated by two-sided Wilcoxon’s signed rank-sum test. The quantification was performed from n = 1,895, 155, and 152 cells for excitatory neurons, Pvalb inhibitory neurons, astrocytes using a subset of spatially separated Xa or Xi in (B), (C), (E), (G), (H) and using a subset of at least one of the Xa or Xi regions (75.3-77.0 Mb) detected in a cell in (D and F) from three biological replicates.

**Fig. S9.**
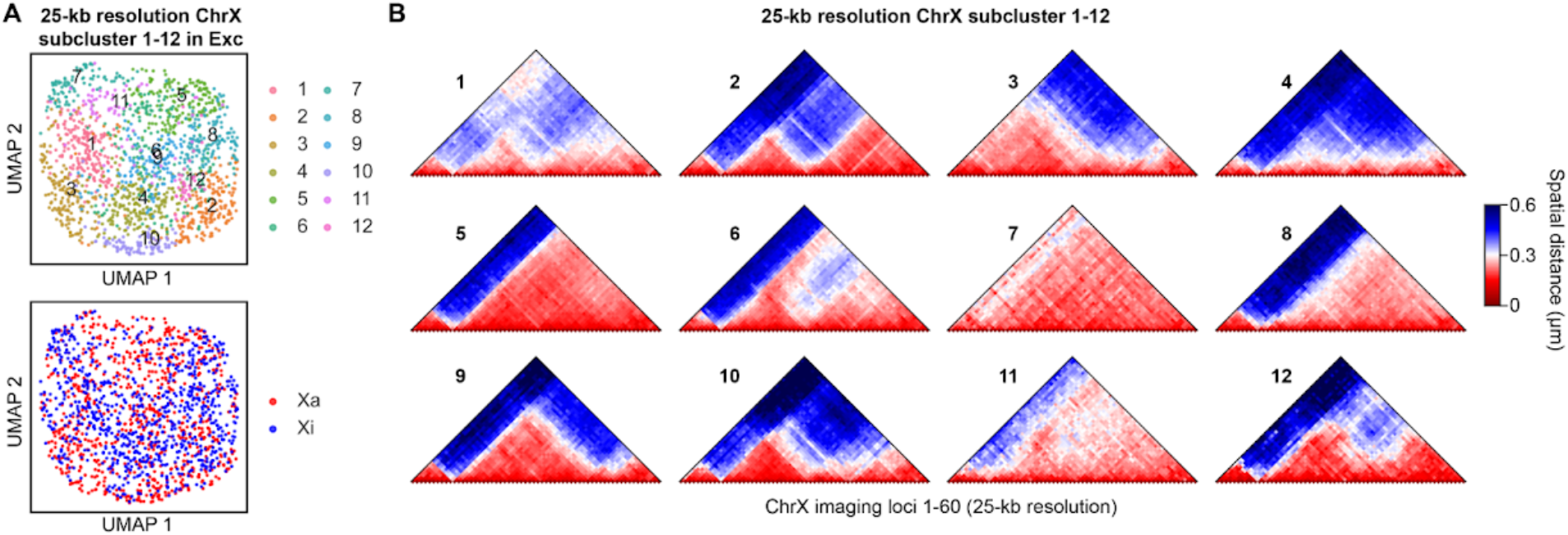
Active and inactive X chromosome organization in subpopulations of excitatory neurons. (A) UMAP of conformation of the 25-kb resolution data for Xa and Xi colored by structural subclusters (top) and Xa and Xi identities (bottom). (B) Pairwise spatial distance maps from all 12 structural clusters of the 25-kb resolution data in chromosome X. n = 805 excitatory neurons with >20% loci detected for both Xa and Xi regions (75.3-77.0 Mb) from three biological replicates in (A) and (B).

## Supplementary Tables

**Table S1** (provided as an Excel file): A list of genomic coordinates for the 3,660 DNA loci in DNA seqFISH+ with a corresponding codebook including unused barcodes.

**Table S2** (provided as an Excel file): A list of target RNAs with corresponding imaging rounds and fluorescent channels used in this study.

**Table S3** (provided as an Excel file): A list of target antibodies and repetitive DNA elements with corresponding imaging rounds and fluorescent channels for each biological replicate.

**Table S4** (provided as a CSV file): A summary of 2,762 cells profiled in this study.

